# LINE-1 ORF2p Expression is Nearly Imperceptible in Human Cancers

**DOI:** 10.1101/744425

**Authors:** Daniel Ardeljan, Xuya Wang, Mehrnoosh Oghbaie, Martin S. Taylor, David Husband, Vikram Deshpande, Jared P. Steranka, Mikhail Gorbounov, Wan Rou Yang, Brandon Sie, H. Benjamin Larman, Hua Jiang, Kelly R. Molloy, Ilya Altukhov, Zhi Li, Wilson McKerrow, David Fenyö, Kathleen H. Burns, John LaCava

## Abstract

**Background:** Long interspersed element-1 (LINE-1, L1) is the major driver of mobile DNA activity in modern humans. When expressed, LINE-1 loci produce bicistronic transcripts encoding two proteins essential for retrotransposition, ORF1p and ORF2p. Many types of human cancers are characterized by L1 promoter hypomethylation, L1 transcription, L1 ORF1p protein expression, and somatic L1 retrotransposition. ORF2p encodes the endonuclease and reverse transcriptase activities required for L1 retrotransposition. Its expression is poorly characterized in human tissues and cell lines.

**Results:** We report mass spectrometry based tumor proteome profiling studies wherein ORF2p eludes detection. To test whether ORF2p could be detected with specific reagents, we developed and validated five rabbit monoclonal antibodies with immunoreactivity for specific epitopes on the protein. These reagents readily detect ectopic ORF2p expressed from bicistronic L1 constructs. However, endogenous ORF2p is not detected in human tumor samples or cell lines by western blot, immunoprecipitation, or immunohistochemistry despite high levels of ORF1p expression. Moreover, we report endogenous ORF1p-associated interactomes, affinity isolated from colorectal cancers, wherein we similarly fail to detect ORF2p. These samples include primary tumors harboring hundreds of somatically-acquired L1 insertions. The new data are available via ProteomeXchange with identifier PXD013743.

**Conclusions:** Although somatic retrotransposition provides unequivocal genetic evidence for the expression of ORF2p in human cancers, we are unable to directly measure its presence using several standard methods. Experimental systems have previously indicated an unequal stoichiometry between ORF1p and ORF2p, but in vivo, the expression of these two proteins may be more strikingly uncoupled. These findings are consistent with observations that ORF2p is not tolerable for cell growth.

## 1 Background

Mobile elements make up nearly half of the human genome [1, 2]. The most prevalent sequences are retrotransposons, which propagate via RNA intermediates, and of these, modern activity resides with the Long INterspersed Element-1 (LINE-1, L1) sequences and those elements mobilized by L1 proteins in trans (reviewed in [3–5]). L1 is the only autonomous (protein-coding), functional retrotransposon in humans, and each of us inherits a distinct complement of active elements [6]. Mobilization occurs after a retrotransposition-competent L1 is transcribed, translated into proteins encoded by its open reading frames (ORFs), ORF1p [7, 8] and ORF2p, and packaged into a ribonucleoprotein (RNP) complexes [9, 10]. The ORF2p encodes an endonuclease [11] that cuts the genomic DNA target site and a reverse transcriptase [12] that generates L1 cDNA.

Many malignant tissues permit L1 expression and somatic retrotransposition [13–23]. Both targeted and genome-wide sequencing efforts have identified thousands of de novo insertions that have occurred across hundreds of human cancers. Several groups have shown that L1 ORF1p expression is a hallmark of many different cancers [18, 24]. Of these many cancer types, it has been shown that L1 upregulation is induced early in the development of ovarian cancers, where ORF1p accumulation is evident within precursor lesions of the fallopian tube [25]. L1 retrotransposition can also contribute directly to cellular transformation; in colon cancers, acquired L1 insertions are known to cause driving mutations in the adenomatous polyposis coli (APC) tumor suppressor [20, 26].

ORF2p is strictly required for retrotransposition, and so its expression in human malignancies can be inferred. Whether the protein can be directly detected has been a matter of some debate. In experimental systems, ORF2p is translated from the bicistronic transcript through an unconventional mechanism [27]. Compared to ORF1p, ectopically expressed ORF2p accumulates in substoichiometric amounts (ORF1p:ORF2p ratio > 30:1) and may be restricted to a subset of cells within a population [9, 10, 28]. Reports of endogenously expressed ORF2p have been more limited than ORF1p [29]. To our knowledge, to date, two groups have independently reported development of monoclonal antibodies recognizing human ORF2p, both using BALB/c mice. One reagent, developed by Belancio and colleagues [30], was reported to provide detection of ectopically-expressed ORF2p only. A second reagent, developed by Sciamanna, Spadafora, and colleagues [31], was reported to detect endogenous ORF2p in several malignant tissues where ORF1p expression has been reported. However, questions have been raised about the specificity of this reagent (see accompanying manuscript by Logan and colleagues). The difficulty directly detecting ORF2p may not simply be a matter of lacking exceptional affinity reagents for sensitive and specific western blotting. As we describe in this report, ORF2p has also eluded robust detection in a systematic, mass spectrometry-based tumor proteome sequencing effort (breast and ovary analyzed here) and in our own immunoprecipitations of ORF1p from resected patient colorectal tumors; this, when ORF1p is robustly detected and captured.

Here we present our perspective on ORF2p detection, including results obtained searching for ORF2p in cancer proteomes as well as probing and analyzing tissue sections and immunoprecipitates. We report endogenous ORF1p-associated interactomes, affinity isolated from colorectal cancers (CRC), in which ORF2p was not found. Moreover, we describe the development of additional reagents to detect ORF2p: 5 rabbit monoclonal antibodies. western blotting, immunoprecipitation, immunohistochemistry, and immunofluorescence demonstrate the utility of these reagents in experimental systems with ectopic LINE-1 expression. However, we cannot detect endogenous ORF2p using these approaches, corroborating previous studies by Belancio [30].

## 2 Results

### 2.1 Detecting ORF1p and ORF2p peptides in tumor mass spectrometry data

We reanalyzed data from Clinical Proteomics Tumor Analysis Consortium (CPTAC) to assess L1 ORF1p and ORF2p protein production in tumors. CPTAC has generated deep mass spectrometry based proteomics data from treatment naive breast [32] and ovarian [33] tumors using isobaric labeling and extensive prefractionation with alkaline reverse phase chromatography followed by inline reverse phase chromatography and Orbitrap mass spectrometry. For the detection of ORF1p and ORF2p peptides, we constructed a protein sequence collection that, in addition to human proteins from Ensembl, also included high confidence LINE-1 protein coding sequences from L1Base2 [34], and used the X! Tandem [35] search engine with the curated databases and the same search parameters as Ruggles et al. [36].

We observed ORF1p in most breast and ovarian tumors (Figure 1A with several peptides observed for the majority of tumors; see Figure 1B and 1C for two examples of quality ORF1 peptide spectrum matches [PSMs]), but there was no clear evidence for ORF2p. Even when we relaxed the filters, the potential evidence for ORF2p peptides was questionable and the majority of PSMs were semi-tryptic and had borderline e-values (≈0.01). We also inspected the potential ORF2p PSMs manually and rejected them because they had several large peaks that could not be explained by fragmentation of the assigned ORF2p peptide. The best ORF2p PSM and only potential evidence for ORF2p that was not rejected is shown in Figure 1D, but this peptide is short and still has two prominent peaks that are not explained by the sequence. In summary, we can reliably observe ORF1p in breast and ovarian tumors using deep mass spectrometry-based proteomics, but, in contrast, the evidence for detection of ORF2p is inconclusive.

**Figure 1:**
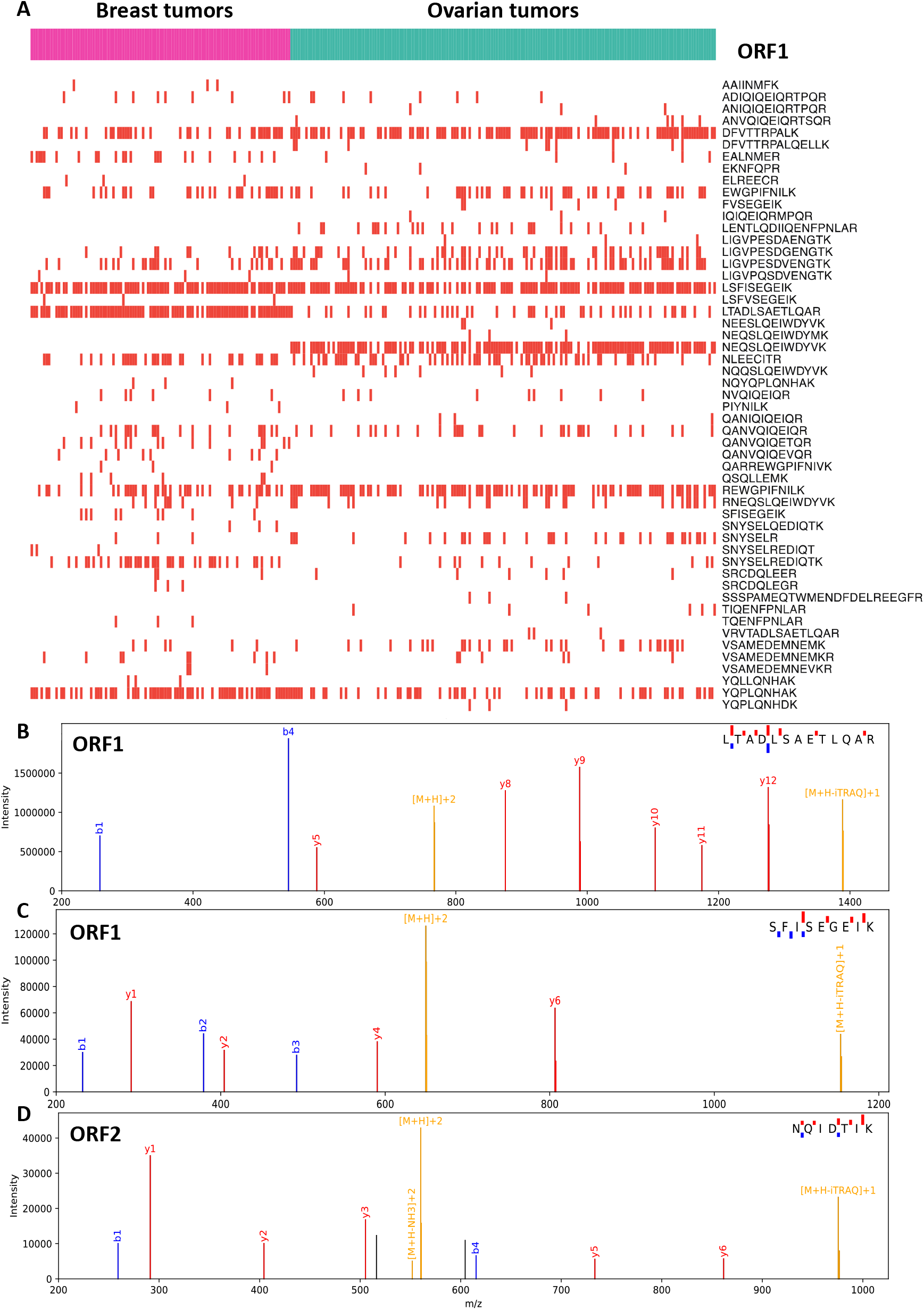
LINE-1 Peptide Detection in Tumor Mass Spectrometry Data. **(A)** ORF1p peptides observed in CPTAC breast and ovarian tumors. Highest quality PSMs that were observed for **(B, C)** ORF1p and **(D)** ORF2p. Precursor ion related peaks are shown in yellow, y-ions in red, b-ions in blue, and unassigned ions in black.

### 2.2 Characterizing L1 immunoprecipitates from CRCs

In our prior work with ectopically expressed L1 RNPs in HEK-293T_*LD*_cells, we readily co-immunoprecipitated ORF1p/ORF2p/*L1* RNA-containing macromolecules, and robustly detected ORF proteins [9, 10]. The detection of ORF2p had, at the time, been a widely recognized problem [29] which we addressed by appending a 3xFlag epitope-tag to the protein. The 3xFlag tag allowed us to robustly capture and detect ORF2p. Because these experiments provided a window on L1 biology only in an ectopic expression context, we wanted to evaluate concordance with pathophysiology. To this end, we sought to isolate L1 RNPs, directly from ORF1p-expressing tumors using an anti-ORF1p affinity medium, comparing and contrasting the results obtained with those of our studies of ectopic L1 expression. We obtained a cohort of CRCs that were shown to be ORF1p positive (+) by immunohistochemistry (IHC) and carried out a preliminary proteomic characterization of three tumors selected from across the ORF1p+ IHC staining spectrum. Figure 2 shows the results we obtained by multiple proteomic methods. Tumor A was the highest ORF1p-expressing case among this group. Tumor C was on the low-to-moderate-end of the expression spectrum and did not yield a distinct, visible ORF1p band after immunoprecipitation (IP): Figure 2A, compare the ORF1p staining intensity in the first (far left, Tumor A), fourth (Tumor B), and eighth (Tumor C) lanes of the gel; **2B** exhibits results obtained with Tumor A using a modified procedure (see Fig. 2 legend and Methods). Also, see Figure 2C for a comparison of ORF1p yield after IP from our highest-expressing ectopic system (pLD401; [10]) to tumors A and B. Although IP from these materials yields ORF1p quantities that are directly comparable, side-by-side western blotting of cell extracts revealed that ectopic expression produced a significantly higher level of ORF1p than these tumors (Fig. 2D). Surprisingly and importantly, ORF2p is only detected on that western blot in the pLD401 ectopic expression positive control. The result did not change when extending the blot exposure time from 2 min (shown) to 30 min (not shown); nor was the result different when using alternative anti-ORF2p antibody clones characterized in this study. Presuming ORF2p has been retrieved from these tumors by co-IP with ORF1p, we conclude the yield is below the lower limit of detection of our blotting, under the conditions tested.

**Figure 2:**
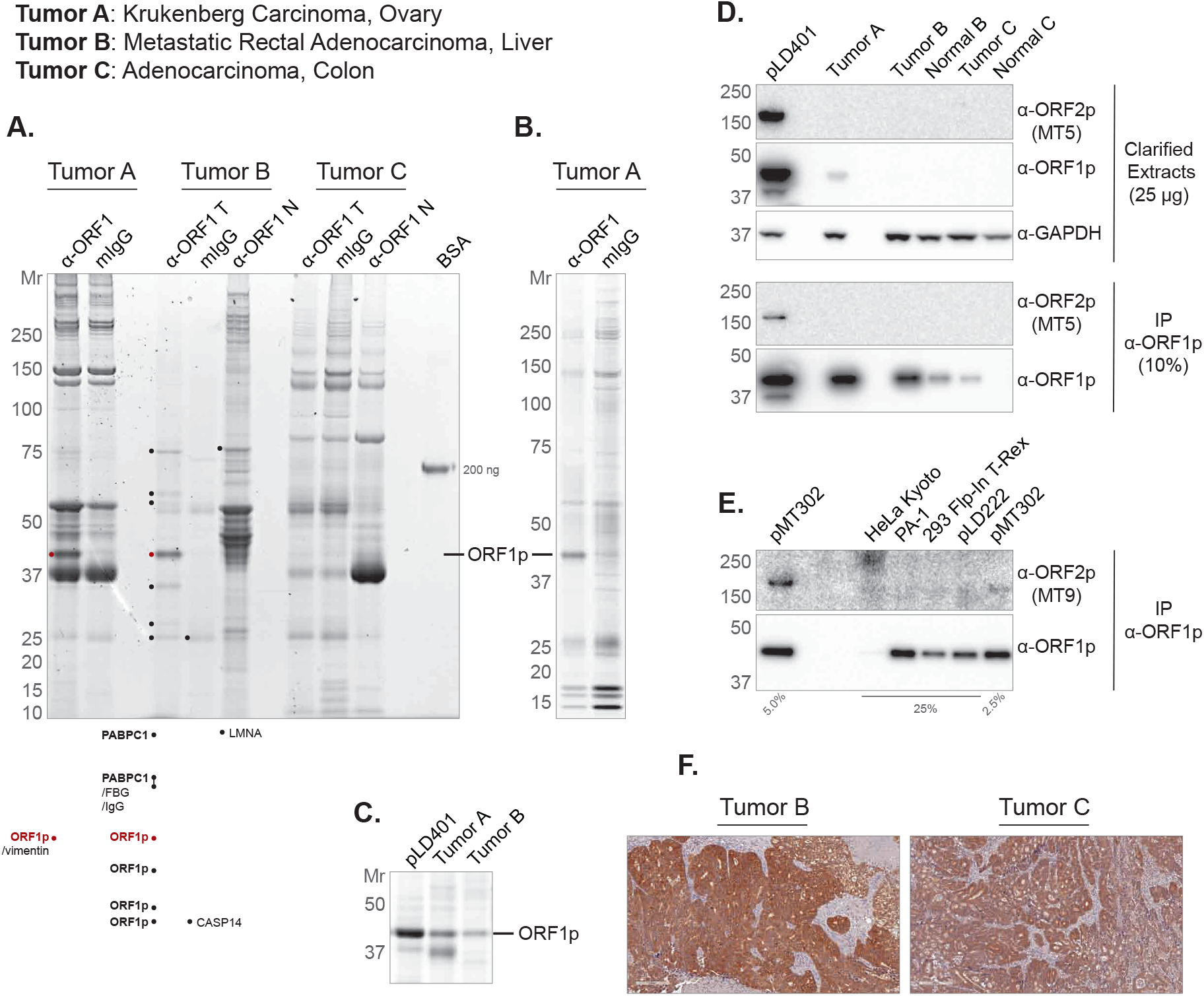
Protein Staining and Western Blotting of anti-ORF1p IPs and extracts. **(A)** Three tumors (labeled at TOP) were used as starting material for ORF1p affinity isolations (*α*-ORF1p), including mock-capture controls using mouse IgG affinity medium with tumor extracts (mIgG), and matched normal tissue with anti-ORF1p affinity medium (*α*-ORF1p N). The eluted material was electrophoresed (4-12% Bis-Tris NuPAGE) and Coomassie G-250 stained [69] and a 200 ng BSA standard is displayed as a staining intensity gauge. Each lane contains a 200 mg-scale isolation using 10 *µ*l of affinity medium. Several bands were cut and analyzed by LC-MS/MS - the highest ranking proteins are listed (see **Methods**) **(B)** Tumor A anti-ORF1p affinity capture was repeated using a slightly modified procedure (see methods). 30% of 100 mg-scale affinity isolations using 15 *µ*l of affinity medium have been electrophoresed and Sypro Ruby stained. **(C)** Comparison of ORF1p yield from anti-ORF1p affinity isolations. pLD401 is a codon-optimized L1 sequence (OrfeusHs), ectopically expressed in HEK-293T_*LD*_[10]. Here, 80% of 100 mg-scale affinity isolations using 10 *µ*l of affinity medium have been electrophoresed and Coomassie G-250 stained. **(D)** Western blotting of the same materials used in (C), including Tumor C and matched normal tissues. Here, 25 *µ*g of the whole cell extract have been probed for ORF2p, ORF1p, and GAPDH as a control. 10% of *α*-ORF1p affinity isolates have also been probed for ORF2p and ORF1p. **(E)** A collection of cell lines were assessed by anti-ORF1p affinity capture. pMT302 is derived from a naturally occurring L1 sequence (L1RP), ectopically expressed in HEK-293T_*LD*_[10]. pLD222 is a plasmid harboring a doxycycline-inducible GFP construct ectopically expressed in HEK-293T_*LD*_; here included as a control for pMT302. **(F)** IHC using *α*-ORF1p on (LEFT) Tumor B and (RIGHT) Tumor C. *α*-ORF2p clones MT5 (panel D) and MT9 (panel E) are described in this study (see Figs. 4 and 5).

To contextualize this result, we developed a similar analysis in a broader selection of cell lines (Fig. 2E). We observed that the yield of endogenous ORF1p by IP was≲ 1/10th the amount observed in HEK-293T_*LD*_expressing L1 ectopically from pMT302. This construct was chosen on account of its milder ectopic expression level. Modified from a naturally occurring L1 sequence (L1RP), expression from pMT302 has been estimated to yield 1/40th the L1 RNA and ORF2p and 1/4th the ORF1p expression typically observed from codon-optimized L1 encoded by pLD401 [10, 29]. PA-1, an ovarian teratocarcinoma cell line known to be permissive for the expression of endogenous L1 [37], stood out among this group - demonstrating 1/10th the ORF1p yield of pMT302 and the highest yield from an endogenous context in this panel. Western blotting also demonstrated ORF1p signal in cell lysates from the panel, but only under probing conditions that increased high mass nonspecific signal in the blot (Fig. S1). Figure 2F shows cognate anti-ORF1p IHC staining results for tumors B and C – corroborating signal intensity difference revealed by the western blots in this figure. ORF2p was not detected, except by co-IP with ORF1p from pMT302.

Believing we exhausted the potential of western blotting for ORF2p detection, we turned to MS-based proteomic analyses. Figure 3 displays the results of a label-free, quantitative MS analysis of affinity captured ORF1p, from the same tumor samples displayed and analyzed in Figure 2 (see also Fig. S2). As expected, we identified L1RE1 (consensus ORF1p) as a significantly enriched protein in each IP set. Taken all together, we observed eight other proteins that we have previously characterized as putative physiological L1 interactors (PABPC1, PABPC4, TUBB, RO60, UPF1, MOV10, HSP90AA1, HSP90AB1); PABPC1/4 being most frequently recovered. We explored the interactors discussed in [38], originating from two studies, conducted by the Moran and Kazazian labs [39, 40]. We observed DHX9 and MATR3 (in Tumor A, set 1), HNRNPC and LARP1 (in Tumor A, set 2), SRSF1 (in Tumor B, set 1 & Tumor B, set 2), SRSF6 and IGF2BP2 (Tumor B, set 2), HNRNPU (in Tumor B, set 2 & Tumor C, set 2), and FAM120A and HNRNPA2B1 (in Tumor C, set 2). Only HNRNPU was observed to be a significant hit in two different patient tumors. Notably, HNRNPU, DHX9, MATR3, HNRNPC, and other RNA binding proteins have been reported to accumulate on L1 and retro-element-derived RNAs; in one hypothesis, insulating these sequences from nuclear RNA processing pathways that might otherwise be deleterious to the retro-element and host genes harboring these sequences [41, 42]. The data can otherwise be summarized as follows: 291 proteins passed at least one t-test (tumor vs. control IP: p-adjusted value of ≤ 0.05 and ≥ two-fold enrichment in tumor IP), 37 passed two, and 22 passed three or more; 21 candidate non-consensus ORF1p loci were detected and of these 12 were observed in both tumor A and tumor B. We observed a candidate phosphorylation site at S18 (156 PSMs from this study). The next most frequent candidate phosphorylation site, S27, received only 31 PSMs; both S18 and S27 phosphosites have previously been reported [43, 44] and have been implicated as (1) functionally important for retrotransposition and (2) mediating an interaction with the peptidyl-prolyl cis/trans isomerase PIN1. The above described findings are further annotated and summarized in **Supplementary Table 1**. Importantly, we did not detect ORF2p in any of these tumor analyses.

**Figure 3:**
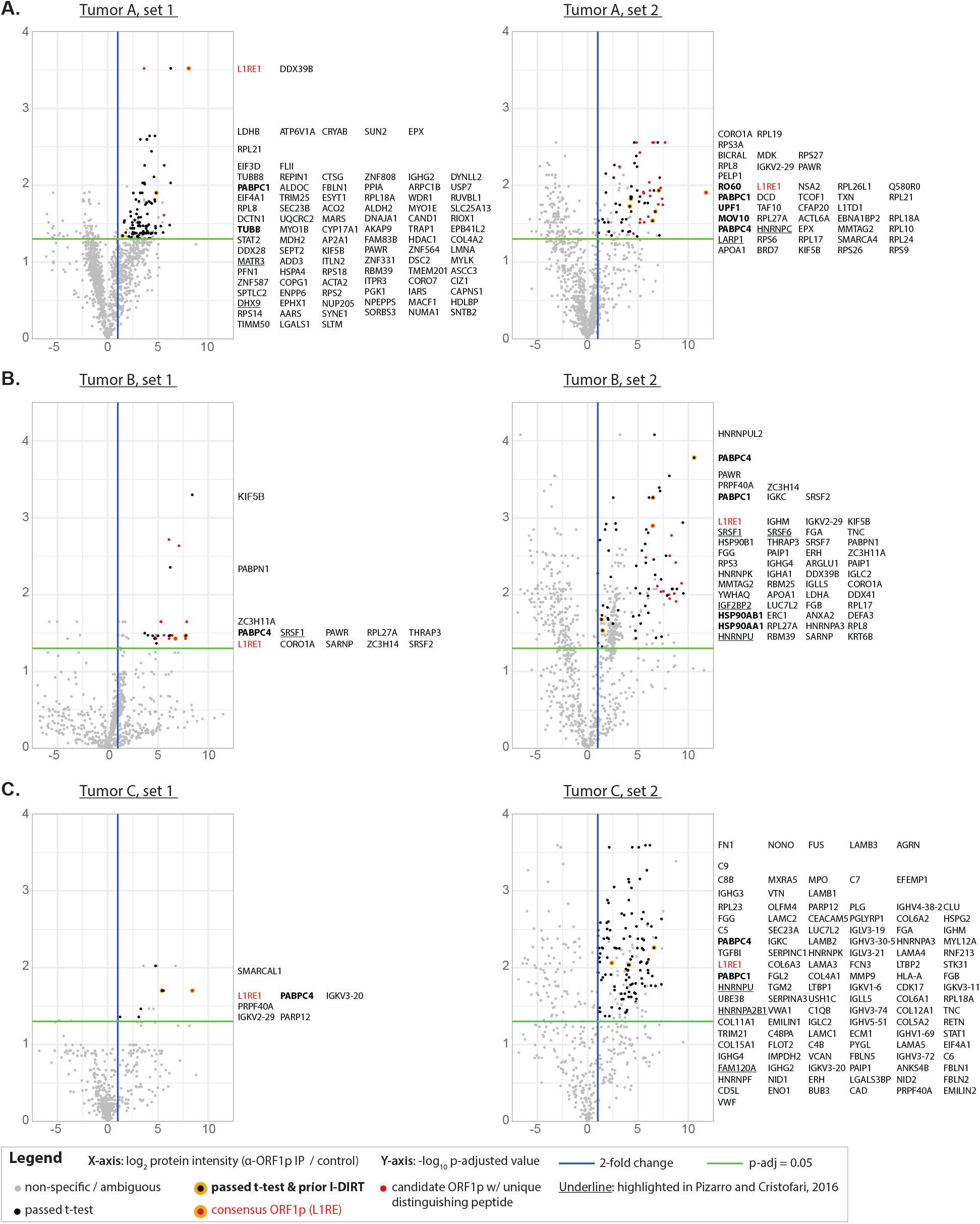
Label-free Quantitative IP-MS analysis. A <u>legend</u> appears at the bottom. Gene symbols corresponding to tumor-specific, quantified proteins displayed at the right of each plot. **(A)** Tumor A, these IPs (set 1 and 2) differ in several experimental parameters (see **Methods**); both sets use a mock IP control (mouse IgG). **(B)** Tumor B, two distinct controls were used: (LEFT) mIgG IP, (RIGHT) matched normal liver, *α*-ORF1p IP. **(C)** Tumor C, controls as for Tumor B. These data correspond to IPs displayed in Figure S2.

### 2.3 Monoclonal antibodies detect human LINE-1 ORF2 protein

We chose the retrotransposition-competent L1RP sequence as an immunogen for generating ORF2p antibodies. Prior to immunization in rabbits, we expressed tagged ORF2 fragments from bacteria, one fragment with the endonu-clease domain (EN, amino acids 1-238, His6 tag) and one fragment containing the reverse transcriptase domain and surrounding sequence (RT, amino acids 238-1061, tagged with mannose binding protein/MBP or a small ubiquitin-like modifier/SUMO) (Fig. 4A). We also expressed full-length flag-tagged ORF2 (ORF2-3xFlag) in Tet-On human embryonic kidney-293T (HEK-293T_*LD*_) cells to screen immune sera. We confirmed fragment purity after Nickel or size-exclusion chromatography (Fig. 4B).

**Figure 4:**
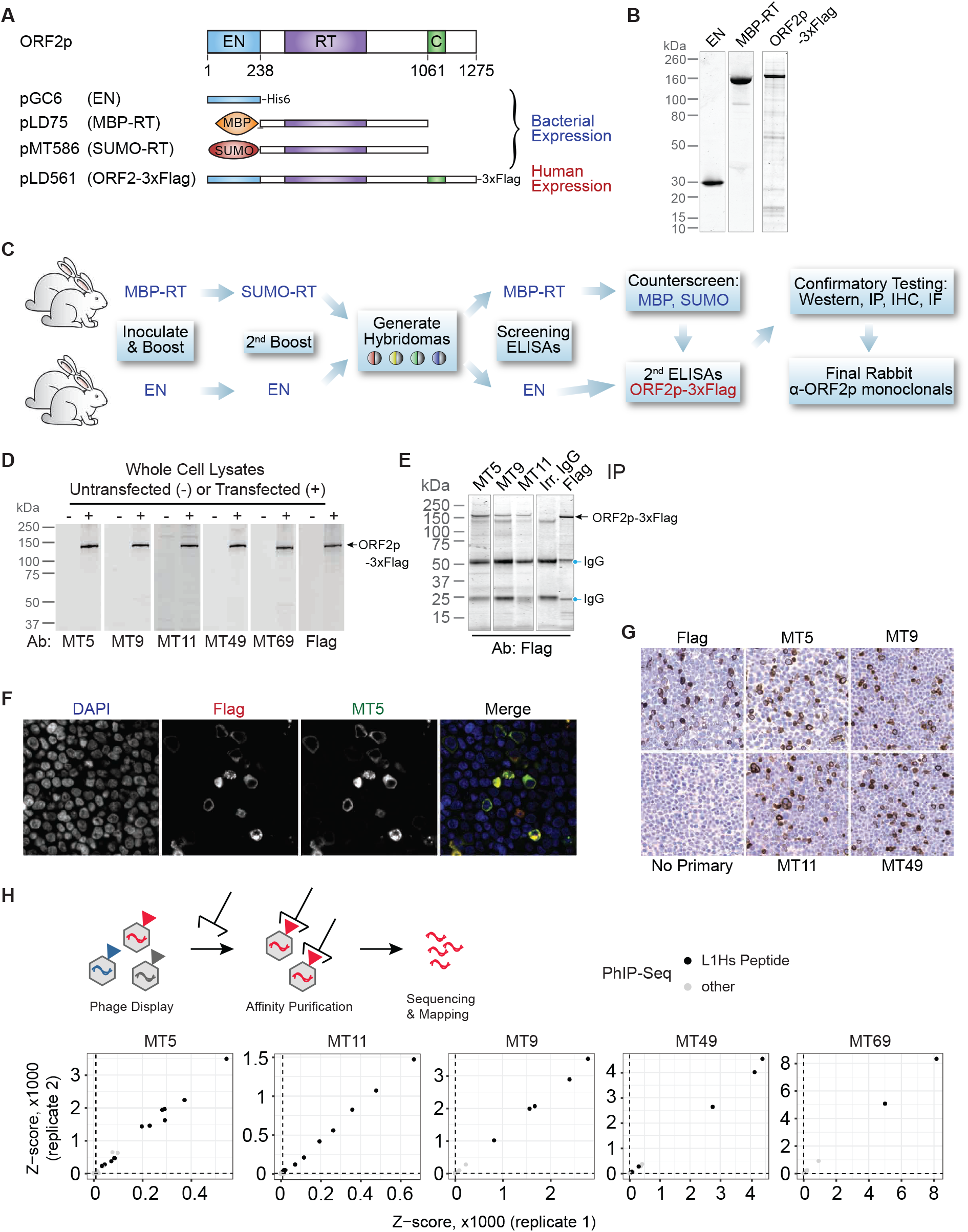
Production of monoclonal ORF2p antibodies. **(A)** Expression constructs used to generate antigens for ORF2p antibody production. **(B)** Coomassie-stained protein electrophoresis gels illustrating purity of ORF2 antigens used in antibody generation. **(C)** Immunization strategy to produce rabbit monoclonal antibodies. **(D)** Western blot detection of ORF2p-3xFlag obtained from HEK-293T_*LD*_cells over-expressing pLD561 (shown in panel A) using 5 different monoclonal antibodies (Ab) compared to anti-Flag. **(E)** Immunoprecipitation of ORF2p-3xFlag using 3 antibodies. **(F)** Immunofluorescence imaging of H EK-293T_*LD*_cells expressing ORF2p-3xFlag showing co-localization with anti-Flag antibody. **(G)** Immunohistochemistry of HEK-293T_*LD*_cells expressing ORF2p-3xFlag with 4 monoclonal antibodies compared to anti-Flag. EN = endonuclease, RT = reverse transcriptase, MBP = mannose binding protein, SUMO = small ubiquitin-like modification. **(H)** Phage-display immunoprecipitation sequencing (PhIP-seq) results. Above, a phage library expresses protein epitopes from the protein-coding genome, which are affinity purified with ORF2 antibodies. DNA sequences are then isolated and sequenced to identify the genes encoding the peptides. Below, results from PhIP-seq using five monoclonal antibodies targeting ORF2p. In each instance, the greatest affinity of the ORF2p monoclonal antibodies is for peptides encoded by L1Hs ORF2p peptides.

For EN-targeting antibodies, we immunized and boosted two rabbits with EN-His6 fragments, then screened hybridoma supernatants by ELISA against purified EN domain and subsequently ORF2-3xFlag. We used the same strategy for RT-targeting antibodies, but used MBP-RT to stimulate the primary immune response and boosted with SUMO-RT to avoid MBP-specific antibody generation. We counter-screened against MBP and SUMO immunoreactivity to identify hybridomas with reactivity for ORF2 (Fig. 4C). Hybridoma supernatants were then tested for their ability to detect ORF2-3xFlag by western blot (Fig. 4D), IP (**4E**), immunofluorescence (**4F**), and IHC (**4G**). We used Flag-antibody as a control to determine whether our ORF2 antibodies detected ORF2-3xFlag. Five monoclonal antibodies (mAbs) were selected based on their ability to detect full-length ORF2-3xFlag by each modality: MT5, MT9, MT11, MT49, and MT69.

### 2.4 ORF2p antibodies are specific for human-specific LINE-1 (L1Hs)

To assess the specificity of antibodies for L1Hs, we employed phage-display immunoprecipitation sequencing (PhIP-seq) [45–47]. We obtained all annotated repeats in the human genome from RepeatMasker and constructed a representational phage-display library which was used in combination with a previously constructed pan human proteome library [48]. The RepeatMasker peptidome was tiled from N-to C-terminus using 56 amino acid peptides with 28 amino acid overlaps. PhIP-Seq enabled us to determine both on- and off-target reactivities of each antibody by sequencing library IPs (Fig. 4H). Each of the five mAbs pulled down several peptides encoded by L1Hs sequences with high significance across both replicates. IP of L1Hs-derived peptides was orders of magnitude more significant compared to any other peptide encoded by the unique and repeat human genome included in the phage displayed libraries. Based on these data, we conclude that these mAbs have minimal off-target reactivity, suggesting that these mAbs should be highly specific to the L1Hs ORF2 protein.

### 2.5 ORF2 antibodies identify non-overlapping epitopes

There are several hundred potentially active L1 loci with intact open reading frames that have been characterized in modern humans [6, 49], and these sequences are highly identical to one another at both the nucleotide and amino acid level. The specific repertoire of L1 loci that is expressed varies among individuals, and by cell or tissue type [20, 22, 50, 51]. To evaluate the potential of our mAbs to detect proteins originating from distinct copies of L1Hs, we mapped the specific epitopes recognized by each antibody and evaluated the conservation of each epitope among L1Hs loci.

We mapped target epitopes using a peptide array of overlapping 15-mers tiling the length of ORF2p. We incubated these arrays with each antibody and then used secondary antibodies conjugated to horseradish peroxidase (HRP) to identify which peptides were identified. Epitopes were identified as the largest contiguous stretch of amino acids that showed mAb binding over background (Fig. 5A). The linear epitopes ranged in length from 6 to 14 amino acids. Each of the 5 epitopes mapped to a discrete, non-overlapping segment of ORF2 in a manner consistent with the purified protein fragments we used for rabbit immunization. The MT49 epitope (DRSTRQ) and MT69 epitope (LHQADLID) occur adjacent to one another and target amino acids on the surface of the endonuclease domain according to a published crystal structure [52]. Both the MT9 epitope (KASRRQEITKIRAE) and MT11 epitope (KELEKQEQT) are located between the annotated EN and RT domains, whereas MT5 identifies an epitope (QDIGVGKD) 300 amino acids from the C terminus, adjacent to the C domain.

**Figure 5:**
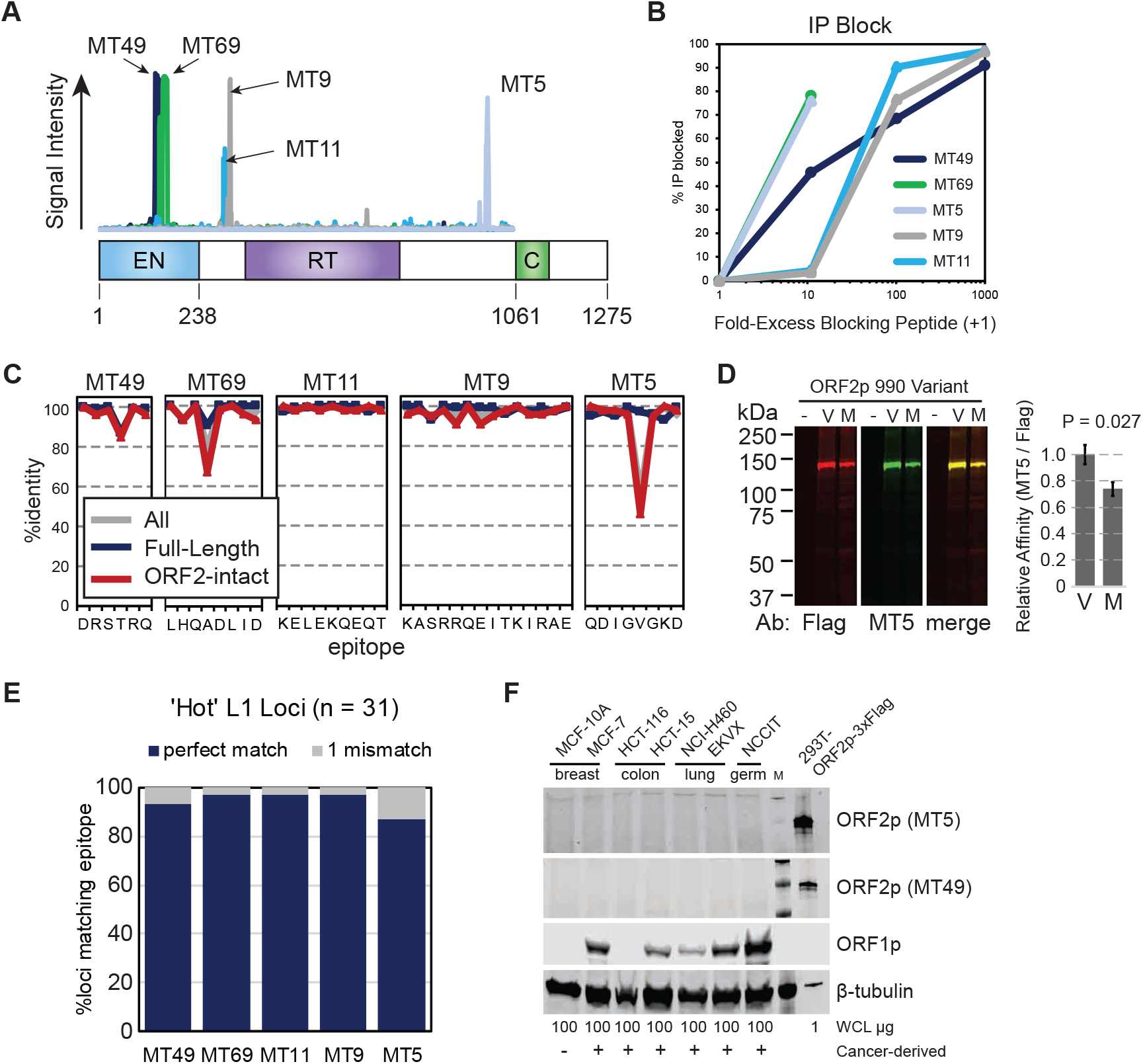
ORF2 mAb can detect endogenous L1Hs. **(A)** Epitopes identified by five ORF2 mAbs are indicated along the linear sequences of the ORF2 protein. **(B)** Immunoprecipitation (IP) blockade of ORF2p pulldown can be achieved by pre-incubating ORF2 mAbs with blocking peptides identified in (A). **(C)** ORF2 mAb epitopes are highly conserved among both full-length and ORF2-intact L1Hs sequences in the human genome. **(D)** Western blot measuring the ability of the MT5 antibody to detect an L1Hs polymorphism at amino acid position 990 reveals that the antibody can detect both alleles. **(E)** Epitope %identities among 31 ‘hot’ or highly active L1Hs sequences as reported by Brouha et al. [49]. **(F)** Western blot of lysates from several ORF1p negative and ORF1p positive cancer cell lines fails to detect ORF2 protein with two different ORF2 mAbs. HEK-293T_*LD*_cells expressing ORF2-3xFlag are included as a positive control.

To validate these epitopes, we pre-incubated mAbs with blocking peptides and attempted IP of ORF2-3xFlag. We found a concentration-dependent blocking activity of each peptide on its corresponding mAb; a range of 10-1000-fold excess peptide was required to achieve this effect depending on the mAb (Fig. 5B). These findings confirm that these epitopes are the antibody targets. PhIP-seq data were also consistent with these being the cognate epitopes recognized by each antibody. Finally, we complimented our finding of antibody specificity by PhIP-seq (Fig. 4H) by performing a BLAST search of these epitopes, which revealed that the only perfect matches in the human genome belong to L1 ORF2p sequences.

### 2.6 ORF2 mAbs are sensitive for many genomic source elements

To evaluate the occurrence of these epitopes in naturally-occuring L1 sequences, we used a census of fixed and commonly-occurring potentially protein-coding L1 elements found in the hg38 reference genome build. We focused on those with intact ORF2 reading frames as previously annotated by L1Base [34, 53]. We performed clustal alignments for two non-overlapping sets of these elements, one consisting of 146 full-length loci (111 L1Hs, 35 L1PA2) and one with 107 ORF2-intact loci. We included consensus sequences of the youngest human-specific L1 (L1Hs) and next-youngest primate-specific L1 (L1PA2) as well as the sequence of L1RP – the antigen against which our mAbs were raised – to compare sequences of the immunogen used for antibody generation against those of other genomic L1 loci. Full-length, intact LINEs are predominantly species-specific L1Hs subfamily, but include some older, primate-specific L1 elements. As expected, full-length and ORF2-intact L1 amino acid sequences are nearly identical for this set (Fig. S3). L1RP-encoded ORF2p is 1,275 amino acids long. Individual, full-length L1 elements had open reading frames that differed from this on average by 16 amino acids (1.25%, range 1-61, Fig. S3), and ORF2-intact L1 loci differed on average by 32 amino acids (2.5%, range 2-79, Fig. S3).

To assess which of the several hundred L1Hs loci could be detected by our mAbs, we used these clustal alignments to evaluate what proportion of L1 loci match each mAb epitopes (Fig. 5C). For each epitope, most full-length L1 loci have amino acid sequences that are 100% identical. The MT11 epitope (KELEKQEQT) and MT9 epitope (KASRRQEITKIRAE) similarly occur nearly universally in ORF2-intact L1 sequences. The greatest discrepancies occurred in the MT5 epitope (QDIGVGKD), where amino acid position 990, which tends to be universal in intact, full-length L1 sequences, is not consistently found in elements selected only for an intact ORF2. Position 990 is typically a valine in L1Hs sequences and a methionine in older elements such as L1PA2 due to a G>A nucleotide substitution (ORF2 position 2,968). We tested whether substituting the L1Hs valine for methionine was sufficient to preclude antibody recognition of the epitope. We created flag-tagged L1RP ORF2p with an M990 substitution, expressed both protein variants in HEK-293T_*LD*_, and performed a western blot using both MT5 and anti-Flag antibodies (Fig. 5D). We detected no signal from MT5 or anti-Flag in untransfected cells. We detected both V990 and M990 variants by both anti-Flag and MT5 antibodies. The M990 variant was detected at a weaker relative intensity by MT5 compared to anti-Flag, indicating a reduced affinity for the ancestral (M990) sequence compared to the derived (V990) sequence. Consequently, we concluded that this single amino acid change within the target epitope may reduce detection sensitivity, but would not prevent detection of L1PA2-encoded ORF2p by this reagent. Among the 31 loci that are considered ‘hot’, or highly active elements [49], mAb epitopes differ by at most 1 amino acid from the L1RP variant (Fig. 5E), suggesting that these youngest L1Hs sequences are likely identifiable by all of the 5 antibodies. Thus, we expect that our mAbs can detect nearly every active L1 encoded in the human genome.

### 2.7 Endogenous ORF2p expression cannot be directly detected in human cancers

Given the lack of ORF2p detection by mass spectrometry of tumor extracts (Fig. 1), we tested for endogenous ORF2p expression in human cancers tissues and cell lines by western blotting, IP-western, IP-MS, and IHC (Figs. 2, 3, and 5). Western blotting and IP-western in several widely used human cell lines, including cells expressing endogenous ORF1p, showed no evidence of ORF2p expression (Figs. 2E and 5F). We conducted IHC in human CRCs with these reagents, including cases known to sustain somatic retrotransposition. ORF1p was readily detectable in all cases evaluated. ORF2p immunostaining, by contrast, showed no consistent signal over isotype controls under standard conditions. Under conditions employing a highly sensitive protocol, immunoreactivity over the isotype control was apparent only inconsistently as cytoplasmic staining with one of the antibodies (MT49, data not shown).

## 3 Discussion

We know that many types of human cancers accumulate somatically-acquired L1 insertions [13-17, 19-21, 23, 54-57]. These insertions have several sequence features that specifically indicate retrotransposition by ORF2p; therefore, we have a high degree of confidence that ORF2p is expressed along with ORF1p in these malignancies. As expected based on the expression level, we reliably detect ORF1p and report a number of co-purifying ORF1p interactors from these cancer samples. Many are proteins that we and others have previously validated. Among them, we noted nuclear RNA binding proteins (e.g. DHX9, HNRNPU, HNRNPC, MATR3) that have been reported by others as L1 interacting proteins, each seen in one tumor sample IP analysis. The significance of these in our ORF1p co-IPs is still debatable, particularly given data from our group and others that ORF1p has limited role and lifespan in the nucleus in cultured cells [9, 28]: do these represent co-assembly of ORF1p with nuclear RNAs (L1 or otherwise); are our ORF1p co-IP populations sampled from cytoplasmic pools of heterogeneous RNAs harboring these proteins [42]; or are these proteins binding ORF1p-containing macromolecules post-lysis (and are therefore artifacts of spurious binding to L1 RNPs in vitro)? Standard LFQ MS control samples, including those used here, are not suited to rule out the latter scenario. Hence, more data are needed in order to dissect their potential significance to L1 molecular physiology.

Despite this strong evidence for ORF2p enzymatic activity, we have demonstrated that endogenous ORF2 protein expression is difficult to reliably directly detect in cancer samples. This is true in the context of: (i.) highly fractionated tumor extracts subjected to mass spectrometry-based shotgun proteomic analysis (as part of the CPTAC initiative); (ii.) anti-ORF1p affinity enrichment followed by western blotting and mass spectrometry; and (iii.) by employing high quality antibodies against ORF2p in standard western blotting, immunoprecipitation, and immunostaining protocols. We also tried to IP ORF2p from CRC tissues and PA-1 cells using anti-ORF2 antibodies, followed by western blotting against ORF2p, but were unable to detect ORF2p (not shown); further optimizations may alter this outcome. While preparing this manuscript, a dataset was released comprising anti-ORF1p co-IPs from H9 human embryonic stem cells with LFQ MS-based analysis [58]; ORF2p detection was not reported. ORF2p peptide detection has previously been reported by MALDI-TOF MS after IP with an anti-ORF2p polyclonal antibody [59]; however, this reagent no longer exists so, further validation and side-by-side comparisons are not possible.

From a bioinformatic perspective the mass spectrometric evidence for ORF1p is strong as several high-quality peptide spectrum matches are observed and their intensities are highly correlated. In contrast, only one ORF2p peptide was observed to have reasonable PSMs (Fig. 1D) but even these have two unexplained medium intensity peaks – increasing the uncertainty that it constitutes an observation of ORF2p. It is also uncomfortable to claim that a protein is observed based on a single relatively short peptide even if it only maps to ORF2p and not to any other protein in the reference human proteome, as it might map to a variant that is present in the sample and not in the reference. In summary, the mass spectrometric evidence for ORF2p in tumors is, at best, questionable. We therefore recommend that ORF2p searches of mass spectrometry data implement additional filters to reduce the likelihood of false positives – these may include the rejection of ORF2p peptide matches when:

1. they exhibit poor fragmentation, particularly where the sequence differs from consensus;
2. the sequence could be explained by deamidation of consensus sequence (D or E in variant corresponds to N or Q in consensus, respectively);
3. the sequence could be explained by a non-tryptic cleavage of consensus sequence; and
4. the peptide matched is not fully tryptic.

Filter 1 is not practical to carry out manually on large data sets but computational approaches are possible. Filters 2, 3, and 4 should be relatively straight forward to execute programmatically in future studies. The difficulty to detect ORF2p has not been missed by the field, yet the enthusiasm to address this challenge may have generated some false starts (we are aware of a newly submitted manuscript from Susan Logan and colleagues re-examining the reliability of the chA1-L1 antibody [31]). Failure to reliably, directly detect endogenous ORF2 protein most likely points to an extremely low steady state abundance for this protein - even in the conditions of L1 de-repression that typify these cancers - in keeping with historical estimates of low ORF2p copy number and high ORF1p:ORF2p stoichiometry [10, 27]. Commonly used ectopic expression systems may not just over-express ORF1p and ORF2p, but may also skew their relative abundances and/or apparent stoichiometry in resulting macromolecules [10]. If this is so, whether these experimental systems change the efficiency of ORF2p translation or overwhelm cellular clearance pathways is unclear. On the other hand, considering the (ectopic) ORF2p western blot signal observed to co-IP with ORF1p from HEK-293T_*LD*_cells, the absolute expression level and lower recovery of ORF1p from a ‘high-L1-expressing tumor’ (tumor A) suggests that the yield is simply too low to expect reliable ORF2p detection, even at the apparent stoichiometry described for ectopic expression in HEK-293T_*LD*_[10]; see Figure 2D: compare ORF1p and ORF2p signals from pLD401 to tumor A.

Moreover, whether there is heterogeneous expression of ORF2p in tumors has as yet not been well addressed. It is possible that relatively rare malignant cells accumulate detectable amounts of ORF2p, and that these have escaped sampling in our IHC experiments. Taylor et al. previously observed that, in the presence of robust ectopic ORF1p expression, only a subset of cells - approximately one-third - also exhibited ORF2p expression [10], indicating a potentially cell-dependent stochastic determinant for ORF2p expression. Similarly, although somatic retrotransposition events result in acquired genomic L1 insertions that are propagated by the clonal expansions of tumors, we do not know whether these accumulate continuously over time or whether instead they reflect discrete, episodic breaches of host defenses against L1. Indeed, there may be active selection against cells with high ORF2p because of cytotoxic effects of the protein or retrotransposition products [60].

## 4. Conclusions

Here, we have evaluated L1 ORF2p expression in human cancers using several independent and orthogonal approaches - one reliant on whole proteome analysis; one leveraging ORF1p interactions to seek evidence of ORF2p; and one employing a series of new, apparently avid and specific monoclonal antibodies for ORF2p detection. While many types of epithelial cancers express levels of ORF1p that are directly detectable by western blotting and mass spectrometry, ORF2p in these cases appears to be only indirectly detectable by gDNA sequencing of de novo L1 insertions. The apparent uncoupling of ORF1p and ORF2p expression is striking, and potentially much more pronounced in vivo than in previously characterized experimental systems. We expect that in the future, more sensitive assays will reveal the now imperceptible quantities of ORF2p. If shotgun proteomic approaches are not sufficient to address ORF2p detection, targeted methods can be developed to maximally leverage the sensitivity of MS instruments (potentially 100s of attomoles [61]). Similarly, approaches such as proximity ligation assays (PLA) or other technologies, may amplify detection by antibodies. Characterizing ORF2p expression and understanding its regulatory mechanisms may have translational importance. If high levels of ORF2p expression are not compatible with malignant cell growth, restoring expression of this protein selectively in cancer cells producing L1 RNA and ORF1p may provide an avenue for therapeutics.

## 5 Methods

### 5.1 Detection of L1 ORF peptides in CPTAC data

The CPTAC discovery breast [32] and ovarian [33] mass spectrometry data was used (available at the CPTAC Data Portal: https://cptac-data-portal.georgetown.edu/cptac/s/S015 and https://cptac-data-portal.georgetown.edu/cptac/s/S020, respectively). For the detection of ORF1p and ORF2p peptides, we constructed a protein sequence collection that, in addition to human proteins from Ensembl, also included high confidence LINE-1 proteins from L1Base2 [34]: 292 ORF1p/ORF2p sequences translated from full-length intact LINE-1 and 107 ORF2p translated from ORF2 intact LINE-1 elements in human, and 89 LINE-1 ORF1p/ORF2p translated from ancestor consensus sequences. In addition, we also included a list of contaminant proteins from the common Repository of Adventitious Proteins (cRAP). We used the X! Tandem [35] (https://www.thegpm.org/tandem/) search engine with the curated databases and the same search parameters as in [36]. In-house scripts were used to parse the X! Tandem outputs to filter for high-quality Peptide-Spectrum Matches (PSMs). Only PSMs that meet the following criteria were retained: the fraction of the intensity of peaks that matched the sequence > 40%, the gaps in the fragmentation were not larger than 3 amino acids, the peptide length >=7 and the e-value <= 0.01. We also eliminated PSMs that match to more than one gene. In order to select a set of reliable peptides from ORF1p, we performed a pair-wise comparison of the peptide quantities only kept the peptides that formed a set that had a Spearman correlation of 0.6 with each other.

### 5.2 Immunoprecipitation

For Figures 2 and 3: handling of cryomilled HEK-293T_*LD*_cells ectopically expressing L1 from pLD401 and pMT302 was previously described [10] [62]. Patient samples were milled and extracted similarly, as previously described [63]. Protein extraction solution: 20 mM HEPES pH 7.4, 500 mM NaCl, 1% (v/v) Triton X-100, 1x Roche Complete EDTA-free protease inhibitors. tumor A was extracted in a separate instance in the same solution with the addition of Promega recombinant RNasin at 1:50 (v:v).

For patient samples subjected to LFQ-MS we used the following parameters: 200 mg-scale, 10 *µ*l of anti-ORF1p (Millipore Sigma #MABC1152) and mouse IgG (Millipore Sigma #I5381) affinity medium were used per 200 mg-scale affinity capture. In addition to the mouse IgG mock affinity capture control, for tumors B and C, we carried out an additional mock affinity capture using the anti-ORF1p antibody and extracts from matched normal tissue, resected at the time the CRC was removed from the patient. Affinity media and clarified extracts were incubated for 1 hr at 4°C, washed three times with extraction solution, and eluted with NuPage sample buffer (Thermo Fisher Scientific #NP0007) at 70°C. After SDS-PAGE (Thermo Fisher Scientific: 1 mm, 4-12% Bis-Tris NuPAGE system), samples were analyzed by general protein staining, western blotting, and/or MS as described in the main text. Samples destined for MS were reduced (DTT) and alkylated (iodoacetamide) prior to electrophoresis. In a second instance, tumor A affinity isolations were conducted at a 100 mg-scale using 15 *µ*l of anti-ORF1p and mouse IgG medium, were extracted and washed (3 × 250 *µ*l washes as opposed to 1 ml) in the presence of 1:50 RNasin (not previously included), and 1x protease inhibitors (normally only present during extraction); approximately 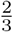 the standard sonication energy was applied (the standard is 15-20 J per 100 mg-scale in a 25% (w:v) extract). In all cases, representative SDS-PAGE lanes are displayed in Figure 2 and the gel plugs used for LFQ-MS are displayed in Figure S2. All panels displayed have been ‘auto tone’ calibrated, respectively, in Adobe Photoshop to maximize the visual contrast across the detected signal range.

For Figures 4 and 5: HEK-293T_*LD*_cells transfected with a plasmid encoding ORF2-3xFlag were lysed by sonication in extraction buffer (50 mM NaCl, 20 mM HEPES pH 7.4, 1% Triton, 1 mM EDTA). 1 *µ*g of each antibody was conjugated to 25 *µ*L of Protein G Dynabeads for 15 minutes at room temperature, then washed with TBST. IP was carried out for 1 hour at room temperature on a rotating wheel with protein lysates diluted by TBST. For peptide blocks, antibodies were pre-incubated with peptide, conjugated to Dynabeads for 15 minutes, and lysate was added for 1 hour at room temperature. After IP, samples were washed in extraction buffer, then eluted from the Dynabeads by heating in LDS (Thermo) at 70C for 10 minutes. Supernatants were then run on Mini TGX gels (Biorad) for western blot with anti-Flag (Sigma) antibody.

### 5.3 Western blotting

For western blots displayed in Figure 2 the following parameters were used: wet transfer (1% [w/v] SDS / 20% [v/v] methanol in transfer buffer) for 90 min / 70 V / 4°C, PVDF membrane (0.45 *µ*m), HRP-conjugated secondary antibodies (see below), and chemiluminescent HRP detection (substrate: Millipore Sigma #WBLUF0100). Blocking was done overnight at 4°C using 5% (w/v) nonfat dry milk in TBST (20 mM Tris-Cl, 137 mM NaCl, 0.1% Tween 20), pH 7.6. Primary antibodies were applied overnight at 4°C in 5% (w/v) BSA in TBST, pH 7.6. Secondary antibodies were applied for 2 hr at room temperature in 5% (w/v) BSA in TBST, pH 7.6. Where appropriate, total protein quantities were estimated using a commercial Bradford reagent. An ImageQuant LAS-4000 system (GE Healthcare) was used for blot imaging on the high sensitivity setting with incremental image capture. ECL signal capture times displayed varied with target from 1 - 5 min and were free of pixel saturation in any signal displayed in the figures. Anti-ORF1p (Millipore Sigma #MABC1152) was used at 0.4 *µ*g/ml; anti-ORF2p (this study) clone MT5 was used at 0.13 *µ*g/ml and clone MT9 was used at 0.71 *µ*g/ml; anti-GAPDH (Cell Signaling #2118) was used at 0.02 *µ*g/ml. Secondary antibodies: anti-mouse HRP conjugate (GE Lifesciences #NV931) and anti-rabbit HRP conjugate (GE Lifesciences #NV934) were used at 1:10,000. All panels displayed have been ‘auto tone’ calibrated, respectively, in Adobe Photoshop to maximize the visual contrast across the detected signal range.

For western blots displayed in Figures 4 and 5, cells were lysed in RIPA buffer, vortexed, and supernatants quantified by BCA. Lysates were reduced in LDS with beta-mercaptoethanol and then polyacrylamide gel electrophoresis was performed on 4-20% Protean Mini TGX gels (Biorad) and transferred to Immobilon PVDF membranes for 15 minutes using mini TGX settings on the Trans-Blot-Turbo system (Biorad). Membranes were incubated with primary antibodies overnight at 4°C (rabbit anti-ORF2 mAbs at 1:1000; mouse anti-Flag M2 (Sigma F1804) at 1:2000), secondary antibodies (all from Licor and used at 1:10,000 dilutions; as appropriate: goat anti-mouse IR680, goat anti-rabbit IR680, goat anti-mouse IR800, goat anti-rabbit IR800) for 1 hour at room temperature, and detection was carried out on the Odyssey Scanner (Licor).

### 5.4 Mass spectrometry

Peptides were resuspended in 10 *µ*L 5% (v/v) methanol, 0.2% (v/v) formic acid and half was loaded onto an EASY-Spray column (Thermo Fisher Scientific, ES800, 15cm × 75*µ*m ID, PepMap C18, 3 *µ*m) via an EASY-nLC 1200 interfaced with a Q Exactive Plus mass spectrometer (Thermo Fisher Scientific). Column temperature was set to 35°C. Using a flow rate of 300 nl/min, peptides were eluted in a gradient of increasing acetonitrile, where Solvent A was 0.1% (v/v) formic acid in water and Solvent B was 0.1% (v/v) formic acid in 95% (v/v) acetonitrile. Peptides were ionized by electrospray at 1.8 – 2.1 kV as they eluted. The elution gradient length was 10 minutes for gel bands and 140 min for all gel plugs except, the second set derived from tumor A, where the gradient length was 190 min. Full scans were acquired in profile mode at 70,000 resolution (at 200 m/z). The top 5 (for gel bands) or 25 (for gel plugs) most intense ions in each full scan were fragmented by HCD. Peptides with charge state 1 or unassigned were excluded. Previously sequenced precursors were also excluded, for 4 s (for gel bands) or 30 s (for gel plugs), within a mass tolerance of 10 ppm. Fragmentation spectra were acquired in centroid mode at 17,500 resolution. The AGC target was 2×105, with a maximum injection time of 200 msec. The normalized collision energy was 24%, and the isolation window was 2 m/z units.

### 5.5 Analysis of excised protein bands and candidate phospho-sites

Proteins labeled in Figure 2A selected for labeling via the following process: The RAW files were converted to MGF format by ProteoWizard [64] and searched against the human protein database with X! Tandem [35], using the following settings: fragment mass error - 10 ppm; parent mass error - 10 ppm; cleavage site - R or K, except when followed by P; maximum missed cleavage sites - 1; maximum valid peptide expectation value - 0.1; fixed modification - carbamidomethylation at C; potential modification - oxidation at M; include reversed sequences - yes. Parameters for the refinement search were: maximum valid expectation value - 0.01; potential modifications - deamidation at N or Q, oxidation or dioxidation at M or W; unanticipated cleavage - yes. For each protein ID list, the proteins were ranked by log E-value; keratins, proteins ranked below trypsin, and non-human proteins were removed; if multiple proteins remained, the nth protein (n>1) was removed if (a) it is homologous to a higher-ranked protein or (b) does not have within 50% the number of PSMs of the top-ranked remaining protein; remaining proteins were listed as IDs for each band. For identification of candidate phosphorylation sites using X! Tandem, the RAW files from tumor IPs (corresponding to Figure 3) were converted to MGF, these were searched against orthogonalized ORF protein sequences (described below), and included the following additional potential modifications during the refinement search: Phospho@S, Phospho@T, Phospho@Y. The best scoring PSMs for phospho S18 and S27 are displayed in **Supplementary Table 1**. The X! Tandem. xml output files are available via ProteomeXchange with identifier PXD013743.

### 5.6 Label-free quantitative analysis

#### Processing RAW data in MaxQuant

we used MaxQuant v1.6.5.0 [65, 66] with default settings and the following adjustments (in brief). Trypsin/P cleavage. Modifications included in protein quantification: Oxidation (M); Acetyl (Protein N-term); Carbamidomethyl (C). Phospho (STY) was searched but excluded from quantification along with unmodified counterpart peptides. Label min. ratio count: 2. Match between runs (within groups of cognate experiments and controls): True. Second peptides: True. Stabilize large LFQ ratios: True. Separate LFQ in parameter groups: False. Require MS/MS for LFQ comparisons: True. We used a protein database composed of the Uniprot human proteome (reviewed), supplemented with non-redundant ORF1p and ORF2p sequences. To increase detection sensitivity, we orthogonalized our ORF1 loci database (from above, Detection of L1 ORF peptides in CPTAC data) within the context of our detected peptides in two steps: (a) retaining loci for which at least one unique peptide was observed and (b) in cases that a peptide was not assigned to any loci in previous step and was commonly shared by several loci, we included only one representative sequence from the group, which was the most different one to consensus ORF1 (L1RE1). The RAW and MaxQuant processed files are available for download via ProteomeXchange with identifier PXD013743.

#### Custom post-processing in R

code can be obtained at https://github.com/moghbaie/L1_CRC_IP_MS.

#### Data preparation

(a) Remove contaminants and reverse protein entries (provided by MaxQuant) and IGHG1. (b) Log2 transformation of intensities (LFQ and iBAQ). (c) Remove proteins with zero values across all cases and controls in a tissue. (d) Impute small values in scenarios that all replicates had zero values for intensity in either cases or controls: we calculated the average (mean) and standard deviation (std) of non-zero values of each replicate and produced small values with the uniform random function between mean – 2*std and mean – 3*std. (e) Impute values for proteins that have zero intensities in one or two replicates in either cases or controls: we built a distribution of deltas from replicates with non-zero protein intensities: 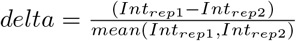; Calculate *µ*_*delta*_, *sd*_*delta*_; Calculate new delta and new Intensity: 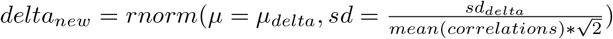; 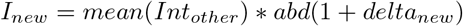

#### Variance Analysis

(a) For tumor A, we performed t-tests between anti-ORF1p IPs and IgG-controls. For tumor B and C, we performed t-tests between tumor and normal tissue after anti-ORF1p IP, as well as between tumor anti-ORF1p IPs and IgG-controls. (b) Adjusted p-values were calculated using Benjamin-Hochberg method. (c) For each entry log2 fold change was calculated between case and control average intensity. (d) Significant proteins from all comparison were integrated (p.adjusted <0.05 & log2fold>1). Because imputation may artifactually inflate the number proteins passing ANOVA, we only accept proteins with MS intensity values in at least two experimental replicates as candidate true positives. Therefore, proteins represented by less than two MS intensity values that passed ANOVA were not labeled in the plots.

### 5.7 Purification of ORF2 Proteins

pGC6, expressing ORF2 endonuclease, is tagged with N-HIS6-TEV. We expressed overnight in bacteria at 16°C, then shifted temperature and induced with IPTG. We froze cell pellets then purified on a nickel column in standard conditions. We cleaved the tag with TEV protease overnight, then performed gel filtration to clean up the untagged protein. pLD75, expressing ORF2 reverse transcriptase tagged with His-MBP, was expressed similarly, purified on a nickel column, then with cation exchange (HiTrap SP FF; GE Healthcare Life Sciences). pLD561, expressing full-length ORF2p-3xFlag, was expressed as a 15 L culture in suspension HEK-293T_*LD*_. We lysed cells with a microfluidizer in 500 mM NaCl buffer with 1% (v/v) Triton X-100, then performed Flag IP using Dynabeads (Thermo Fisher Scientific) coupled to anti-Flag M2 (Millipore Sigma) followed by 3xFlag elution.

### 5.8 Generation of monoclonal antibodies

Rabbit monoclonal antibodies were developed with Abcam (Cambridge, MA). For EN-targeting antibodies, rabbits were immunized and boosted with EN, screened by ELISA for EN affinity, and then hybridoma supernatants were tested against ORF2-3xFlag by ELISA. For RT-targeting antibodies, rabbits were immunized with MBP-tagged RT, boosted with SUMO-tagged RT, then screened by ELISA with MBP-RT and counter-screened with MBP and SUMO to eliminate clones that were specific for MBP or SUMO.

### 5.9 ORF2p induction in HEK-293T_*LD*_cells

Plasmid DNA was miniprepped using the Zyppy Miniprep Plasmid DNA kit (Zymo, Irvine, CA) or PureLink HiPure Plasmid Midiprep Kit (Thermo Fisher, Waltham, MA). These were transfected into Tet-On HEK-293T_*LD*_cells [10] by incubating 3 *µ*g plasmid DNA with 9 *µ*L Fugene HD (Promega, Madison, WI) in 100 *µ*L Optimem for 15 min, then adding dropwise to 6-well plates containing 500,000 cells per well. 1 *µ*g/ml doxycycline was added at the time of transfection and cells were then used for immunoprecipitation, immunofluorescence, immunohistochemistry, or western blot assays 24 hr later.

### 5.10 ORF1p and ORF2p Immunohistochemistry

For cells, HEK-293T_*LD*_cells expressing a plasmid encoding ORF2-3xFlag were admixed with untransfected HEK-293T_*LD*_and pelleted, fixed in 10% formalin for 24 hr, then processed into paraffin-embedded blocks. For human tissue samples, de-identified paraffin-embedded blocks were obtained from the Pathology Department at Massachusetts General Hospital. Formalin-fixed paraffin embedded tissues were sectioned at 5 *µ*m onto glass slides, heated to 65°C for 20 min, and then rehydrated by serial washes in xylene, ethanol (100% / 90% / 75%), and water. IHC was performed with the DAKO EnVision+ System-HRP kit (cat# K4006, Agilent, Santa Clara, CA). Antigen retrieval was performed using Target Retrieval Solution for 20 minutes at > 90°C, then slides were blocked with peroxidase block and then 2% (w/v) BSA in PBS. Primary antibody incubation with ORF1p was performed at 1:5000 for 1 hr at room temperature and with ORF2 T49 ov ernight at 4°C at a final concentration of 10 *µ*g/ml, and secondary HRP mouse polymer secondaries were used to label primary antibody with chromogen upon DAB addition. Hematoxylin was used as a nuclear counterstain. Slides were then dehydrated in serial washes and coverslips were placed. Scoring was performed by a trained pathologist.

### 5.11 Immunofluorescence (IF)

HEK-293T_*LD*_cells expressing a plasmid encoding ORF2-3xFlag were admixed with untransfected HEK-293T_*LD*_and pelleted, fixed in 10% formalin for 24 hr, then processed into paraffin-embedded blocks. IF was performed on 5*µ*M sections. Slides were processed as for IHC but using AlexaFlour-conjugated secondary antibodies (anti-rabbit 488 and anti-mouse 555). Imaging was performed on a Zeiss Confocal Microscope.

### 5.12 Antibody Epitope Mapping

A library of peptide based epitope mimics was synthesized using solid-phase Fmoc synthesis. An amino functionalized polypropylene support was obtained by grafting with a proprietary hydrophilic polymer formulation, followed by reaction with t-butyloxycarbonyl-hexamethylenediamine (BocHMDA) using dicyclohexylcarbodiimide (DCC) with N-hydroxybenzotriazole (HOBt) and subsequent cleavage of the Boc-groups using trifluoroacetic acid (TFA). Standard Fmoc-peptide synthesis was used to synthesize peptides on the amino-functionalized solid support by custom modified JANUS liquid handling stations (Perkin Elmer). The binding of antibody to each of the synthesized peptides was tested in a pepscan-based ELISA. The peptide arrays were incubated with primary antibody solution (overnight at 4°C). After washing, the peptide arrays were incubated with a 1:1000 dilution of anti-rabbit IgG HRP conjugate (DAKO) for one hour at 25°C. After washing, the peroxidase substrate 2,2’-azino-di-3-ethylbenzthiazoline sulfonate (ABTS) and 20 *µ*l/ml of 3 percent H2O2 were added. After one hour, the color development was measured with a charge coupled device (CCD) - camera and an image processing system. Epitope targets were read as the largest contiguous stretch of amino acids shared by all peptides recognized by the primary antibodies.

### 5.13 Peptide blocking experiments

Blocking peptides were chosen to span two extra amino acids and were N-terminally acetylated and C-terminally amidated. Peptides were resuspended in acetic acid or ammonium acetate depending on their charge characteristics. To pre-block antibodies, 10, 100, or 1000-times excess peptide by weight was incubated with primary antibody mixture on a rotating wheel at 4°C overnight. The next day, pre-blocked or no-block antibodies were conjugated to protein G dynabeads for 15 min at room temperature, then ORF2p-3xFlag HEK-293T_*LD*_lysate was added for 1 hour at room temperature. The remaining protocol is the same as described above for immunoprecipitation.

### 5.14 Phage immunoprecipitation and DNA sequencing

The PhIP-Seq assay was described previously [47]. Approximately 100 ng of each mAb was added to the combined T7 bacteriophage human peptidome library (unique genome and repetitive element sublibrary addition, 1 x 105 plaque forming units for each phage clone in each library) and incubated with rotation overnight at 4°C in deep 96-well plates in 1 mL total volume of phosphate-buffered saline. Negative controls for data normalization included eight mock immunoprecipitation reactions on each plate. mAb-phage complexes were captured by magnetic beads (20*µ*L of protein A-coated and 20 *µ*L of protein G-coated, catalog numbers 10002D and 10004D, Invitrogen, Carlsbad, CA) for 4 hours at 4°C with rotation and processed using the Agilent Bravo liquid handling system (Agilent Technologies, Santa Clara, CA). Beads were washed twice with 0.1% NP-40 in Tris-buffered saline (50 mM Tris-HCl with 150mM NaCl, pH 7.5), resuspended in 20 *µ*L of a Herculase II-containing PCR mix (catalog number 600679, Agilent Technologies), and ran for 20 PCR cycles followed by a second 20-cycle PCR using 2 *µ*L of the initial PCR products to add barcodes and P5/P7 Illumina sequencing adapters. Pooled PCR products were sequenced using an Illumina HiSeq 2500 (Illumina, San Diego, CA) in rapid mode (50 cycles, single end reads). Data were normalized and analyzed using a z-scores algorithm according to Yuan et. al. [67].

## 6 Ethics approval and consent to participate

Formalin fixed paraffin embedded (FFPE) and fresh frozen CRC samples were collected at Massachusetts General Hospital Department of Pathology as de-identified patient samples in accordance with Exemption 4 of research involving human subjects from the NIH.

## 7 Availability of data and materials

*Proteomics data*: The mass spectrometry proteomics data have been deposited to the ProteomeX-change Consortium via the PRIDE [68] partner repository with the dataset identifier PXD013743. *R code*: https://github.com/moghbaie/L1_CRC_IP_MS

## 10 Authors’ contributions

D.A., D.F, K.H.B., and J.L. wrote and revised the manuscript. J.L., M.S.T., and V.D. conceived of the tumor anti-ORF1p affinity proteomic analyses; J.L. designed the experimental approach. M.S.T. and K.H.B. developed the strategy for generating ORF2p antibodies. M.S.T. and D.H. isolated the recombinant protein fragments used as immunogens. D.A., M.S.T., J.P.S., and K.H.B. conducted experiments and interpreted data for selecting clones and characterizing the antibodies. M.S.T and V.D. collected and curated the clinical samples used in the immunohistochemistry and immunoprecipitation experiments. Mi.G. conducted immunohistochemistry experiments. H.B.L. led the group conducting PhIP-seq characterizations of these reagents: W.R.Y collected and annotated the repeat sequences to be included in the PhIP library, B.S. performed the cloning to generate the library. H.J. conducted the immunoprecipitations, acquired SDS-PAGE/protein staining and western blotting data, and conducted sample workup for mass spectrometry. K.R.M acquired MS data. MS data processing, analyses, and visualization were carried out by J.L., K.R.M., I.A., X.W., Z.L., W.M., D.F., and M.O. LINE-1 protein sequences database was constructed by X.W. Custom R code for LFQ-MS written by I.A. and M.O. Biological interpretation and drafting of text related to L1 detection in CPTAC data was done by D.F. and J.L. Biological interpretation of the results and drafting of the text related to affinity proteomic analyses was done by J.L. Drafting of the text related to ORF2p antibodies was done by D.A. and K.H.B. All authors read and approved the final manuscript.

## 11 Acknowledgements

We thank Profs. Brian T. Chait and Michael P. Rout for providing NCDIR resources and research infrastructure, used in part to facilitate this study. We thank Leila Saba for contributing to the second immunoprecipitation experiment of tumor A, as well as Svetlana Kalmykova and Maria Pashkova for contributing to our imputation approach. We also thank Mark Grivainis who examined sequence variant alignments between L1 RNAs and proteins, Uri Laserson who developed and implemented the pepsyn software used in this project, and Reema Sharma for technical assistance selecting ORF2p antibody clones.

---

**Figure S1:**
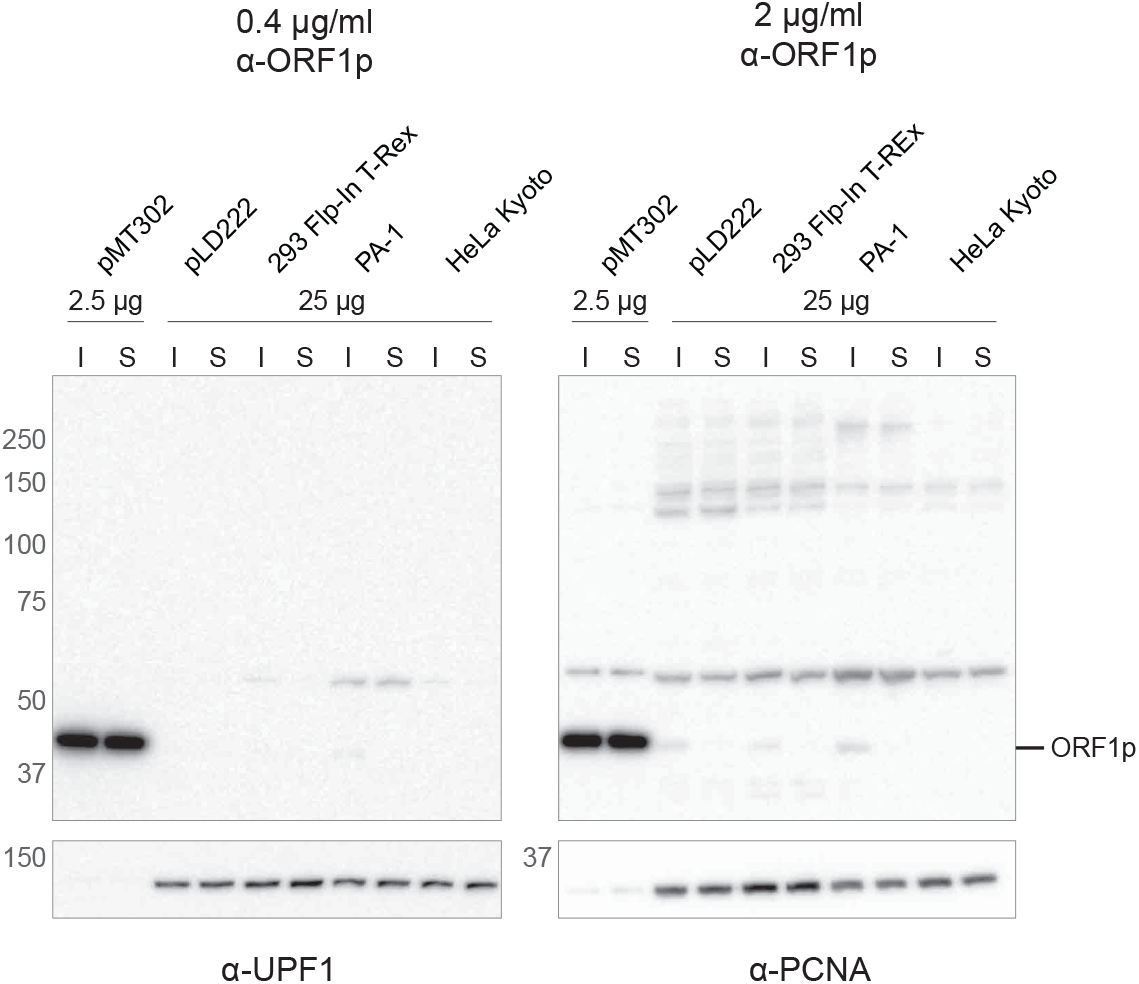
Western blot α-ORF1p titer to detect endogenous ORF1p in clarified cell extracts. The concentration of *α*-ORF1p used is given along the top; the source of each cell extract is given below that, and each accords to Figure 2E. The quantity of clarified cell extracts used, in *µ*g total protein, follows below each extract source. **I**: clarified extract used as an input for *α*-ORF1p affinity capture; **S**: immuno-depleted extracts after incubation with *α*-ORF1p affinity medium. (**Left blot image**) 1x *α*-ORF1p concentration - ORF1p signal is observed in with ectopic expression (pMT302) and at just above background in PA-1. *α*-UPF1 provided as a loading control (NYU1.1B6, 1:1000 [70]). (**Right blot image**) 5x *α*-ORF1p concentration - ORF1p signal is observed in all cases except HeLa Kyoto. An increase in non-specific signal is also observed elsewhere on the blot. *α*-PCNA is provided as a loading control (Santa Cruz Biotechnology, Inc. #sc-56; 1:1000).

**Figure S2:**
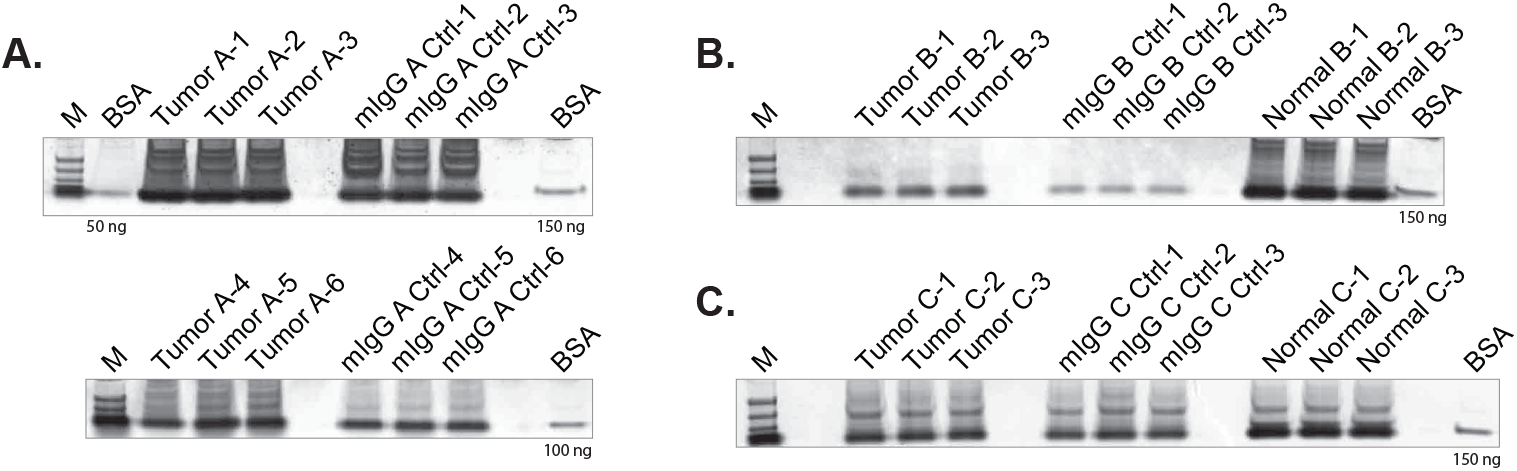
Coomassie G-250 stained gel plugs used for in-gel digestion followed by MS. A panel is shown for every replicate included in the LFQ-MS analysis. **(A)** Tumor A (Krukenberg Carcinoma, Ovary) was subjected to two independent affinity isolations with different parameters (see **Methods**). Each isolation included three replicates using *α*-ORF1p-coupled affinity medium to capture ORF1p from the tumor extracts (Tumor A-1 to A-6), and three replicates using mouse IgG-coupled affinity medium to sample non-specific background from the same extracts (mIgG A Ctrl-1 to Ctrl-6). **(B)** Tumor B (Metastatic Rectal Adenocarcinoma, Liver): including three replicates using *α*-ORF1p-coupled affinity medium to capture ORF1p from the tumor extracts (Tumor B-1 to B-3), three replicates using mouse IgG-coupled affinity medium to sample non-specific background from the same extracts (mIgG B Ctrl-1 to Ctrl-6), and three replicates using anti-ORF1p-coupled affinity medium to capture ORF1p from matched normal tissue extracts (Normal B-1 to B-3). **(C)** Tumor C (Adenocarcinoma, Colon): including three replicates using *α*-ORF1p-coupled affinity medium to capture ORF1p from the tumor extracts (Tumor C-1 to C-3), three replicates using mouse IgG-coupled affinity medium to sample non-specific background from the same extracts (mIgG C Ctrl-1 to Ctrl-6), and three replicates using *α*-ORF1p-coupled affinity medium to capture ORF1p from matched normal tissue extracts (Normal C-1 to C-3).

**Figure S3:**
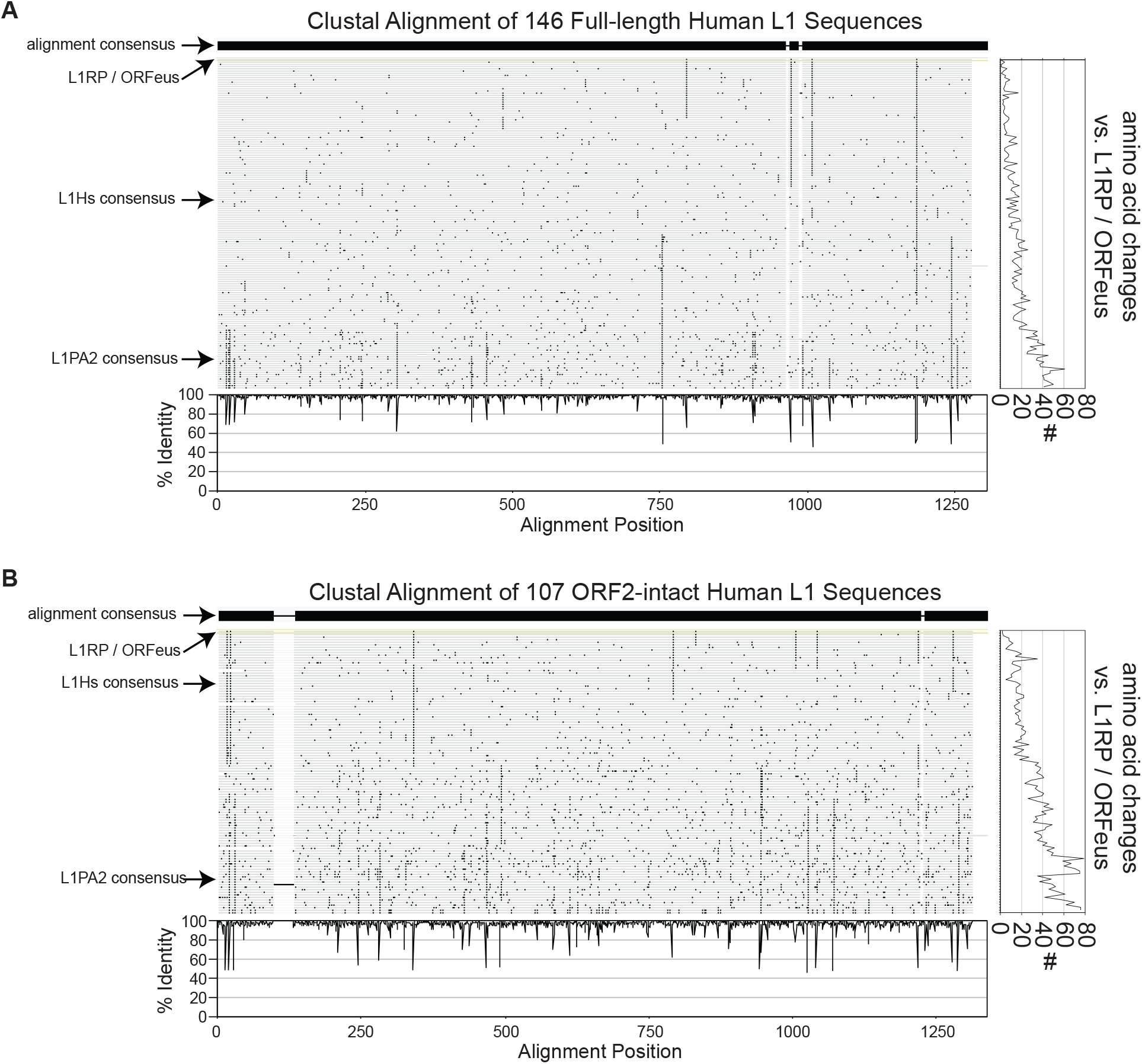
CLUSTAL Alignments of ORF2 protein sequences. ORF2p protein sequences were obtained from the L1Base database of reference L1Hs sequences. **(A)** Alignment of 146 full-length L1 sequences. **(B)** Alignment of 107 ORF2-intact L1 sequences. In the center tiles, black bars indicate amino acid positions where the L1 in that row differs from the CLUSTAL alignment consensus sequence. The ‘% agreement,’ or identity, at each amino acid position is quantified below the center tiles. On the right, the number of amino acid changes of a particular L1 compared to the immunogen, L1RP, is quantified.

## References

1. Lander ES, Linton LM, Birren B, Nusbaum C, Zody MC, Baldwin J, Devon K, Dewar K, Doyle M, FitzHugh W, et al: Initial sequencing and analysis of the human genome. Nature 2001, 409:860–921.

2. RepeatMasker Open-4.0. [http://www.repeatmasker.org]

3. Huang CR, Burns KH, Boeke JD: Active transposition in genomes. Annu Rev Genet 2012, 46:651–675.

4. Kazazian HH, Jr., Moran JV: The impact of L1 retrotransposons on the human genome. Nat Genet 1998, 19:19–24.

5. Kazazian HH, Jr., Moran JV: Mobile DNA in Health and Disease. N Engl J Med 2017, 377:361–370.

6. Beck CR, Collier P, Macfarlane C, Malig M, Kidd JM, Eichler EE, Badge RM, Moran JV: LINE-1 retrotransposition activity in human genomes. Cell 2010, 141:1159–1170.

7. Martin SL, Branciforte D, Keller D, Bain DL: Trimeric structure for an essential protein in L1 retrotrans-position. Proc Natl A cad Sci U S A 2003, 100:13815–13820.

8. Khazina E, Truffault V, Buttner R, Schmidt S, Coles M, Weichenrieder O: Trimeric structure and flexibility of the L1ORF1 protein in human L1 retrotransposition. Nat Struct Mol Biol 2011, 18:1006–1014.

9. Taylor MS, Altukhov I, Molloy KR, Mita P, Jiang H, Adney EM, Wudzinska A, Badri S, Ischenko D, Eng G, et al: Dissection of affinity captured LINE-1 macromolecular complexes. Elife 2018, 7.

10. Taylor MS, LaCava J, Mita P, Molloy KR, Huang CR, Li D, Adney EM, Jiang H, Burns KH, Chait BT, et al: Affinity proteomics reveals human host factors implicated in discrete stages of LINE-1 retrotransposition. Cell 2013, 155:1034–1048.

11. Feng Q, Moran JV, Kazazian HH, Jr., Boeke JD: Human L1 retrotransposon encodes a conserved en-donuclease required for retrotransposition. Cell 1996, 87:905–916.

12. Mathias SL, Scott AF, Kazazian HH, Jr., Boeke JD, Gabriel A: Reverse transcriptase encoded by a human transposable element. Science 1991, 254:1808–1810.

13. Burns KH: Transposable elements in cancer. Nat Rev Cancer 2017, 17:415–424.

14. Tang Z, Steranka JP, Ma S, Grivainis M, Rodic N, Huang CR, Shih IM, Wang TL, Boeke JD, Fenyo D, Burns KH: Human transposon insertion profiling: Analysis, visualization and identification of somatic LINE-1 insertions in ovarian cancer. Proc Natl Acad Sci U S A 2017, 114:E733–E740.

15. Doucet-O’Hare TT, Sharma R, Rodic N, Anders RA, Burns KH, Kazazian HH, Jr.: Somatically Acquired LINE-1 Insertions in Normal Esophagus Undergo Clonal Expansion in Esophageal Squamous Cell Carcinoma. Hum Mutat 2016, 37:942–954.

16. Doucet-O’Hare TT, Rodic N, Sharma R, Darbari I, Abril G, Choi JA, Young Ahn J, Cheng Y, Anders RA, Burns KH, et al: LINE-1 expression and retrotransposition in Barrett’s esophagus and esophageal carcinoma. Proc Natl Acad Sci U S A 2015, 112:E4894–4900.

17. Rodic N, Steranka JP, Makohon-Moore A, Moyer A, Shen P, Sharma R, Kohutek ZA, Huang CR, Ahn D, Mita P, et al: Retrotransposon insertions in the clonal evolution of pancreatic ductal adenocarcinoma. Nat Med 2015, 21:1060–1064.

18. Rodic N, Sharma R, Sharma R, Zampella J, Dai L, Taylor MS, Hruban RH, Iacobuzio-Donahue CA, Maitra A, Torbenson MS, et al: Long interspersed element-1 protein expression is a hallmark of many human cancers. Am J Pathol 2014, 184:1280–1286.

19. Iskow RC, McCabe MT, Mills RE, Torene S, Pittard WS, Neuwald AF, Van Meir EG, Vertino PM, Devine SE: Natural mutagenesis of human genomes by endogenous retrotransposons. Cell 2010, 141:1253–1261.

20. Scott EC, Gardner EJ, Masood A, Chuang NT, Vertino PM, Devine SE: A hot L1 retrotransposon evades somatic repression and initiates human colorectal cancer. Genome Res 2016, 26:745–755.

21. Lee E, Iskow R, Yang L, Gokcumen O, Haseley P, Luquette LJ, 3rd, Lohr JG, Harris CC, Ding L, Wilson RK, et al: Landscape of somatic retrotransposition in human cancers. Science 2012, 337:967–971.

22. Tubio JM, Li Y, Ju YS, Martincorena I, Cooke SL, Tojo M, Gundem G, Pipinikas CP, Zamora J, Raine K, et al: Mobile DNA in cancer. Extensive transduction of nonrepetitive DNA mediated by L1 retrotransposition in cancer genomes. Science 2014, 345:1251343.

23. Helman E, Lawrence MS, Stewart C, Sougnez C, Getz G, Meyerson M: Somatic retrotransposition in human cancer revealed by whole-genome and exome sequencing. Genome Res 2014, 24:1053–1063.

24. Ardeljan D, Taylor MS, Ting DT, Burns KH: The Human Long Interspersed Element-1 Retrotransposon: An Emerging Biomarker of Neoplasia. Clin Chem 2017.

25. Pisanic TR, 2nd, Asaka S, Lin SF, Yen TT, Sun H, Bahadirli-Talbott A, Wang TH, Burns KH, Wang TL, Shih IM: Long Interspersed Nuclear Element 1 Retrotransposons Become Deregulated during the Development of Ovarian Cancer Precursor Lesions. Am J Pathol 2019, 189:513–520.

26. Miki Y, Nishisho I, Horii A, Miyoshi Y, Utsunomiya J, Kinzler KW, Vogelstein B, Nakamura Y: Disruption of the APC gene by a retrotransposal insertion of L1 sequence in a colon cancer. Cancer Res 1992, 52:643–645.

27. Alisch RS, Garcia-Perez JL, Muotri AR, Gage FH, Moran JV: Unconventional translation of mammalian LINE-1 retrotransposons. Genes Dev 2006, 20:210–224.

28. Mita P, Wudzinska A, Sun X, Andrade J, Nayak S, Kahler DJ, Badri S, LaCava J, Ueberheide B, Yun CY, et al: LINE-1 protein localization and functional dynamics during the cell cycle. Elife 2018, 7.

29. Dai L, LaCava J, Taylor MS, Boeke JD: Expression and detection of LINE-1 ORF-encoded proteins. Mob Genet Elements 2014, 4:e29319.

30. Sokolowski M, DeFreece CB, Servant G, Kines KJ, deHaro DL, Belancio VP: Development of a monoclonal antibody specific to the endonuclease domain of the human LINE-1 ORF2 protein. Mob DNA 2014, 5:29.

31. De Luca C, Guadagni F, Sinibaldi-Vallebona P, Sentinelli S, Gallucci M, Hoffmann A, Schumann GG, Spadafora C, Sciamanna I: Enhanced expression of LINE-1-encoded ORF2 protein in early stages of colon and prostate transformation. Oncotarget 2016, 7:4048–4061.

32. Mertins P, Mani DR, Ruggles KV, Gillette MA, Clauser KR, Wang P, Wang X, Qiao JW, Cao S, Petralia F, et al: Proteogenomics connects somatic mutations to signalling in breast cancer. Nature 2016, 534:55–62.

33. Zhang H, Liu T, Zhang Z, Payne SH, Zhang B, McDermott JE, Zhou JY, Petyuk VA, Chen L, Ray D, et al. Integrated Proteogenomic Characterization of Human High-Grade Serous Ovarian Cancer. Cell 2016, 166:755–765.

34. Penzkofer T, Jager M, Figlerowicz M, Badge R, Mundlos S, Robinson PN, Zemojtel T: L1Base 2: more retrotransposition-active LINE-1s, more mammalian genomes. Nucleic Acids Res 2017, 45:D68–D73.

35. Craig R, Beavis RC: TANDEM: matching proteins with tandem mass spectra. Bioinformatics 2004, 20:1466–1467.

36. Ruggles KV, Tang Z, Wang X, Grover H, Askenazi M, Teubl J, Cao S, McLellan MD, Clauser KR, Tabb DL, et al: An Analysis of the Sensitivity of Proteogenomic Mapping of Somatic Mutations and Novel Splicing Events in Cancer. Mol Cell Proteomics 2016, 15:1060–1071.

37. Garcia-Perez JL, Morell M, Scheys JO, Kulpa DA, Morell S, Carter CC, Hammer GD, Collins KL, O’Shea KS, Menendez P, Moran JV: Epigenetic silencing of engineered L1 retrotransposition events in human embryonic carcinoma cells. Nature 2010, 466:769–773.

38. Pizarro JG, Cristofari G: Post-Transcriptional Control of LINE-1 Retrotransposition by Cellular Host Factors in Somatic Cells. Front Cell Dev Biol 2016, 4:14.

39. Moldovan JB, Moran JV: The Zinc-Finger Antiviral Protein ZAP Inhibits LINE and Alu Retrotransposition. PLoS Genet 2015, 11:e1005121.

40. Goodier JL, Cheung LE, Kazazian HH, Jr.: Mapping the LINE1 ORF1 protein interactome reveals associated inhibitors of human retrotransposition. Nucleic Acids Res 2013, 41:7401–7419.

41. Warkocki Z, Krawczyk PS, Adamska D, Bijata K, Garcia-Perez JL, Dziembowski A: Uridylation by TUT4/7 Restricts Retrotransposition of Human LINE-1s. Cell 2018, 174:1537–1548 e1529.

42. Attig J, Agostini F, Gooding C, Chakrabarti AM, Singh A, Haberman N, Zagalak JA, Emmett W, Smith CWJ, Luscombe NM, Ule J: Heteromeric RNP Assembly at LINEs Controls Lineage-Specific RNA Processing. Cell 2018, 174:1067–1081 e1017.

43. Rigbolt KT, Prokhorova TA, Akimov V, Henningsen J, Johansen PT, Kratchmarova I, Kassem M, Mann M, Olsen JV, Blagoev B: System-wide temporal characterization of the proteome and phosphoproteome of human embryonic stem cell differentiation. Sci Signal 2011, 4:rs3.

44. Cook PR, Jones CE, Furano AV: Phosphorylation of ORF1p is required for L1 retrotransposition. Proc Natl Acad Sci U S A 2015, 112:4298–4303.

45. Larman HB, Zhao Z, Laserson U, Li MZ, Ciccia A, Gakidis MA, Church GM, Kesari S, Leproust EM, Solimini NL, Elledge SJ: Autoantigen discovery with a synthetic human peptidome. Nat Biotechnol 2011, 29:535–541.

46. Larman HB, Laserson U, Querol L, Verhaeghen K, Solimini NL, Xu GJ, Klarenbeek PL, Church GM, Hafler DA, Plenge RM, et al: PhIP-Seq characterization of autoantibodies from patients with multiple sclerosis, type 1 diabetes and rheumatoid arthritis. J Autoimmun 2013, 43:1–9.

47. Mohan D, Wansley DL, Sie BM, Noon MS, Baer AN, Laserson U, Larman HB: PhIP-Seq characterization of serum antibodies using oligonucleotide-encoded peptidomes. Nat Protoc 2018, 13:1958–1978.

48. Xu GJ, Shah AA, Li MZ, Xu Q, Rosen A, Casciola-Rosen L, Elledge SJ: Systematic autoantigen analysis identifies a distinct subtype of scleroderma with coincident cancer. Proc Natl Acad Sci U S A 2016, 113:E7526–E7534.

49. Brouha B, Schustak J, Badge RM, Lutz-Prigge S, Farley AH, Moran JV, Kazazian HH, Jr.: Hot L1s account for the bulk of retrotransposition in the human population. Proc Natl Acad Sci U S A 2003, 100:5280–5285.

50. Philippe C, Vargas-Landin DB, Doucet AJ, van Essen D, Vera-Otarola J, Kuciak M, Corbin A, Nigumann P, Cristofari G: Activation of individual L1 retrotransposon instances is restricted to cell-type dependent permissive loci. Elife 2016, 5.

51. Deininger P, Morales ME, White TB, Baddoo M, Hedges DJ, Servant G, Srivastav S, Smither ME, Concha M, DeHaro DL, et al: A comprehensive approach to expression of L1 loci. Nucleic Acids Res 2016.

52. Weichenrieder O, Repanas K, Perrakis A: Crystal structure of the targeting endonuclease of the human LINE-1 retrotransposon. Structure 2004, 12:975–986.

53. Penzkofer T, Dandekar T, Zemojtel T: L1Base: from functional annotation to prediction of active LINE-1 elements. Nucleic Acids Res 2005, 33:D498–500.

54. Ewing AD, Gacita A, Wood LD, Ma F, Xing D, Kim MS, Manda SS, Abril G, Pereira G, Makohon-Moore A, et al: Widespread somatic L1 retrotransposition occurs early during gastrointestinal cancer evolution. Genome Res 2015, 25:1536–1545.

55. Solyom S, Ewing AD, Rahrmann EP, Doucet T, Nelson HH, Burns MB, Harris RS, Sigmon DF, Casella A, Erlanger B, et al. Extensive somatic L1 retrotransposition in colorectal tumors. Genome Res 2012, 22:2328–2338.

56. Nguyen THM, Carreira PE, Sanchez-Luque FJ, Schauer SN, Fagg AC, Richardson SR, Davies CM, Jesuadian JS, Kempen MHC, Troskie RL, et al: L1 Retrotransposon Heterogeneity in Ovarian Tumor Cell Evolution. Cell Rep 2018, 23:3730–3740.

57. Shukla R, Upton KR, Munoz-Lopez M, Gerhardt DJ, Fisher ME, Nguyen T, Brennan PM, Baillie JK, Collino A, Ghisletti S, et al: Endogenous retrotransposition activates oncogenic pathways in hepatocellular carcinoma. Cell 2013, 153:101–111.

58. Vuong LM, Pan S, Donovan PJ: Proteome Profile of Endogenous Retrotransposon-Associated Complexes in Human Embryonic Stem Cells. Proteomics 2019, 19:e1900169.

59. Ergun S, Buschmann C, Heukeshoven J, Dammann K, Schnieders F, Lauke H, Chalajour F, Kilic N, Stratling WH, Schumann GG: Cell type-specific expression of LINE-1 open reading frames 1 and 2 in fetal and adult human tissues. J Biol Chem 2004, 279:27753–27763.

60. Wallace NA, Belancio VP, Deininger PL: L1 mobile element expression causes multiple types of toxicity. Gene 2008, 419:75–81.

61. Faktor J, Sucha R, Paralova V, Liu Y, Bouchal P: Comparison of targeted proteomics approaches for detecting and quantifying proteins derived from human cancer tissues. Proteomics 2017, 17.

62. Taylor MS, LaCava J, Dai L, Mita P, Burns KH, Rout MP, Boeke JD: Characterization of L1-Ribonucleoprotein Particles. Methods Mol Biol 2016, 1400:311–338.

63. LaCava J, Jiang H, Rout MP: Protein Complex Affinity Capture from Cryomilled Mammalian Cells. J Vis Exp 2016.

64. Chambers MC, Maclean B, Burke R, Amodei D, Ruderman DL, Neumann S, Gatto L, Fischer B, Pratt B, Egertson J, et al: A cross-platform toolkit for mass spectrometry and proteomics. Nat Biotechnol 2012, 30:918–920.

65. Cox J, Mann M: MaxQuant enables high peptide identification rates, individualized p.p.b.-range mass accuracies and proteome-wide protein quantification. Nat Biotechnol 2008, 26:1367–1372.

66. Tyanova S, Temu T, Cox J: The MaxQuant computational platform for mass spectrometry-based shot-gun proteomics. Nat Protoc 2016, 11:2301–2319.

67. Yuan T, Mohan D, Laserson U, Ruczinski I, Baer AN, Larman HB: Improved Analysis of Phage ImmunoPrecipitation Sequencing (PhIP-Seq) Data Using a Z-score Algorithm. bioRxiv 2018:285916.

68. Perez-Riverol Y, Csordas A, Bai J, Bernal-Llinares M, Hewapathirana S, Kundu DJ, Inuganti A, Griss J, Mayer G, Eisenacher M, et al: The PRIDE database and related tools and resources in 2019: improving support for quantification data. Nucleic Acids Res 2019, 47:D442–D450.

69. Candiano G, Bruschi M, Musante L, Santucci L, Ghiggeri GM, Carnemolla B, Orecchia P, Zardi L, Righetti PG: Blue silver: a very sensitive colloidal Coomassie G-250 staining for proteome analysis. Electrophoresis 2004, 25:1327–1333.

70. Mita P, Lhakhang T, Li D, Eichinger DJ, Fenyo D, Boeke JD: Fluorescence ImmunoPrecipitation (FLIP): a Novel Assay for High-Throughput IP. Biol Proced Online 2016, 18:16.

